# Generic and context-dependent gene modulations during *Hydra* whole body regeneration

**DOI:** 10.1101/587147

**Authors:** Yvan Wenger, Wanda Buzgariu, Chrystelle Perruchoud, Gregory Loichot, Brigitte Galliot

## Abstract

The cnidarian *Hydra* is a classical model of whole-body regeneration. Historically, *Hydra* apical regeneration has received more attention than its basal counterpart, most studies considering these two regenerative processes independently. We present here a transcriptome-wide comparative analysis of apical and basal regeneration after decapitation and mid-gastric bisection, augmented with a characterization of positional and cell-type expression patterns in non-regenerating animals. The profiles of 25’637 *Hydra* transcripts are available on HydrATLAS (https://hydratlas.unige.ch), a web interface allowing a convenient access to each transcript profile. These data indicate that generic impulse-type modulations occur during the first four hours post-amputation, consistent with a similar integration of injury-related cues on both sides of the amputation plane. Initial divergences in gene regulations are observed in regenerating tips between four and eight hours post-amputation, followed by a dramatic transcriptomic reprogramming between eight and 16 hours when regulations become sustained. As expected, central components of apical patterning, *Wnt3* and *HyBra1*, are among the earliest genes up-regulated during apical regeneration. During early basal regeneration, a BMP signaling ligand (*BMP5-8c*) and a potential BMP inhibitor (*NBL1)* are up-regulated, suggesting that BMP signaling is involved in the basal organizer, as supported by higher levels of phosphorylated Smad in the basal region and by the LiCl-induced extension of *NBL1* expression. By contrast, upon ectopic activation of Wnt/β-catenin signaling, *NBL1* is no longer expressed, basal differentiation is not maintained and basal regeneration is abolished. A tight cross-talk between Wnt/β-catenin apically and BMP signaling basally appears necessary for maintaining and regenerating *Hydra* anatomy.

## Introduction

Spectacular examples of regeneration are found in each phylum of the animal kingdom. Nevertheless, regeneration only occurs in some species scattered across the tree of life, even, closely related species exhibit very different regenerative abilities (Brockes and Kumar, 2008; Bely and Nyberg, 2010). Furthermore, even in organisms considered to have a high regenerative potential, efficient regeneration is typically restricted to specific organs or body parts. It is commonly accepted that a fundamental requirement for regeneration is the capacity of affected tissues to undergo patterning. The initiation of patterning upon injury, and likely patterning itself, involve shared (injury-related) and context-specific (identity-related) gene regulations that drive the specification of appropriate body parts. However, our understanding on these common and specific molecular aspects of regeneration is incomplete. Why some animals have high regenerative abilities and others do not remains an open question that fosters important promises in regenerative medicine. Indeed, answering this question could lead to the implementation of strategies to unlock the regenerative potential of species with limited regenerative abilities like mammals.

The freshwater cnidarian *Hydra* has been a classical model for the study of regeneration since it was shown that any piece of the *Hydra* body column is able to regenerate complete animals (Trembley, 1744; Galliot et al., 2006). *Hydra* consists in a tubular, ~0.5-1 cm long gastric cavity terminated at the apical extremity by a single opening named mouth, surrounded by a dome-shaped structure named hypostome and a ring of tentacles, and at the basal extremity by a peduncle that surmounts the basal disc (**Fig. S1A**). *Hydra* tissues host three stem cells populations, which are located in the central body column and are not interconvertible. The two distinct unipotent epithelial stem cells populate the inner gastrodermis (gESC) and the outer epidermis (eESC), while the multipotent interstitial stem cells (ISCs) are spread along the epidermis, giving rise to sensory-motor and ganglia neurons, mechano-sensory cells named nematocytes (or cnidocytes), gland cells, and finally germ cells when the animals undergo gametogenesis (Bosch, 2009). All three stem cell populations actively self-renew along the central body column while the extremities predominantly contain terminally differentiated cells that progressively get sloughed off.

Upon bisection, *Hydra* regenerate any missing part of their body within two to three days (Fig. S1). In fact, both extremities of the adult animal host a developmental organizer that actively maintain the terminal structures in intact animals (Browne, 1909; MacWilliams, 1983a; Broun and Bode, 2002; Vogg et al., 2016). Upon bisection, such organizers rapidly form, an apical organizer at the tip of the lower part, and a basal organizer at the tip of the upper part. ESCs carry the morphogenetic potential, but both the continuous homeostatic patterning process and the injury-induced regeneration rely on the cross-talk between interstitial derivatives and ESCs (Kobatake and Sugiyama, 1989; Chera et al., 2009).

How injury can promote different molecular and cellular responses in head- and foot-regenerating tips that lead to the reactivation of specific developmental programs remains largely unknown. Biochemical and immunological analyses showed that upon mid-gastric bisection the immediate phosphorylation of RSK and the subsequent modulation of the CREB DNA-binding complex are necessary for head regeneration, indicating that as in bilaterians (Cordeiro and Jacinto, 2013), transcriptional-independent modulations play a key role in the injury response (Galliot et al., 1995; Kaloulis et al., 2004; Wenger, 2014). Indeed, the immediate activation of the MAPK/ERK pathway is necessary for injury-induced cell death (Chera et al., 2009; Chera et al., 2011). A number of genes, required for head or foot regeneration, are up-regulated in the regenerating tips at the immediate, early or early-late phases of the regenerative processes (see in (Galliot et al., 2006). Among them, some are putative target genes of the MAPK/ERK pathway, as *Wnt3*, which is required for *de novo* head patterning and plays a central role in head regeneration (Guder et al., 2006; Lengfeld et al., 2009; Nakamura et al., 2011). Concerning foot regeneration, not much is known on the transition from wound healing to basal patterning. The NK2 transcription factor appears as a key regulator of basal patterning (Grens et al., 1996), possibly directing the expression of the epithelia-peptide pedibin (Hoffmeister, 1996; Grens et al., 1999; Thomsen et al., 2004).

In bilaterians, unbiased transcriptomics/proteomics approaches have proven useful to determine the genetic programs involved in the regeneration of complex structures, such as body parts in planarians (Kao et al., 2013) or *Hydra* (Petersen et al., 2015), appendages in zebrafish, lizards, salamanders (Hutchins et al., 2014; Bryant et al., 2017; Rabinowitz et al., 2017), heart in zebrafish (Lai et al., 2017) and mammals (Quaife-Ryan et al., 2017), lens in salamanders (Looso et al., 2013). These programs drive wound healing, stem cell recruitment and/or somatic tissue re-specification and 3D-patterning, the latter including the re-activation of specific developmental patterning mechanisms. However, the precise outline between wound healing and morphogenesis have been challenging to trace because of several confounding factors in comparative approaches such as large differences between the regenerative strategies of specific structures within a given organism (heart tissue, limb, lens), the developmental/maturation stage of the regenerating organisms and the occurrence of species-specific innovations (Sanchez-Alvarado and Tsonis, 2006; Galliot and Ghila, 2010; Tanaka and Reddien 2011).

To circumvent these limitations, we took advantage of the ability of *Hydra* to undergo a rapid wound healing on each side of the bisection plane followed by a whole-body regeneration through two distinct programs, apical from the lower part and basal from the upper part. We performed a systematic quantitative transcriptomic analysis of the gene regulations during apical and basal regeneration. In addition, we mapped through the same genome-wide approach the expression profiles in intact animals (i) along the body axis, (ii) in each stem cell population, (iii) in the epithelial stem cells once the interstitial stem cells have been eliminated. Results are made available in a Hydra gene expression atlas (<u>HydrATLAS</u>: hydrATLAS.unige.ch), a web interface based on the blast alignment tool (Fig. S2). HydrATLAS provides with an easy and convenient access to sequences from several H. vulgaris strains and visual representations of transcript levels relevant to regional specification during homeostasis and regeneration, cellular plasticity.

## Results

### Global assessment of spatial expression patterns during homeostasis

To gain insights into prominent gene expression patterns along *Hydra* body axis, we applied a strategy based on RNA-seq. Pieces of tissues were dissected from five regions along its main axis, each piece being about 250-500 µm long: the most apical region was obtained after decapitation immediately below the tentacles, the adjacent region (R1) corresponded to the upper body column, two adjacent slices (R3, R4) were dissected from the central body column after mid-gastric bisection, the fifth region included a portion of the peduncle and the basal disk (see Fig. 1A). To reduce noise and improve the reproducibility of the study, we restricted the analyses presented here to transcripts with at least 100 reads in one or more sample, representing 25’637 transcripts. Among these, we found 8’678 transcripts (34% of the dataset) that exhibit statistically significant variations along the body column as deduced from the Likelihood ratio test with a false discovery rate (FDR) < 10^−3^ and variations larger than 10% between the maximum and the minimum values. By grouping these homeostatic patterns in 12 clusters (termed hom-01 to hom-12), we identify a large number of transcripts differentially regulated at the apical and basal poles compared to the three regions of the body column where expression values appear rather homogeneous (Fig. 1B, **Table S1**).

**Figure 1.**
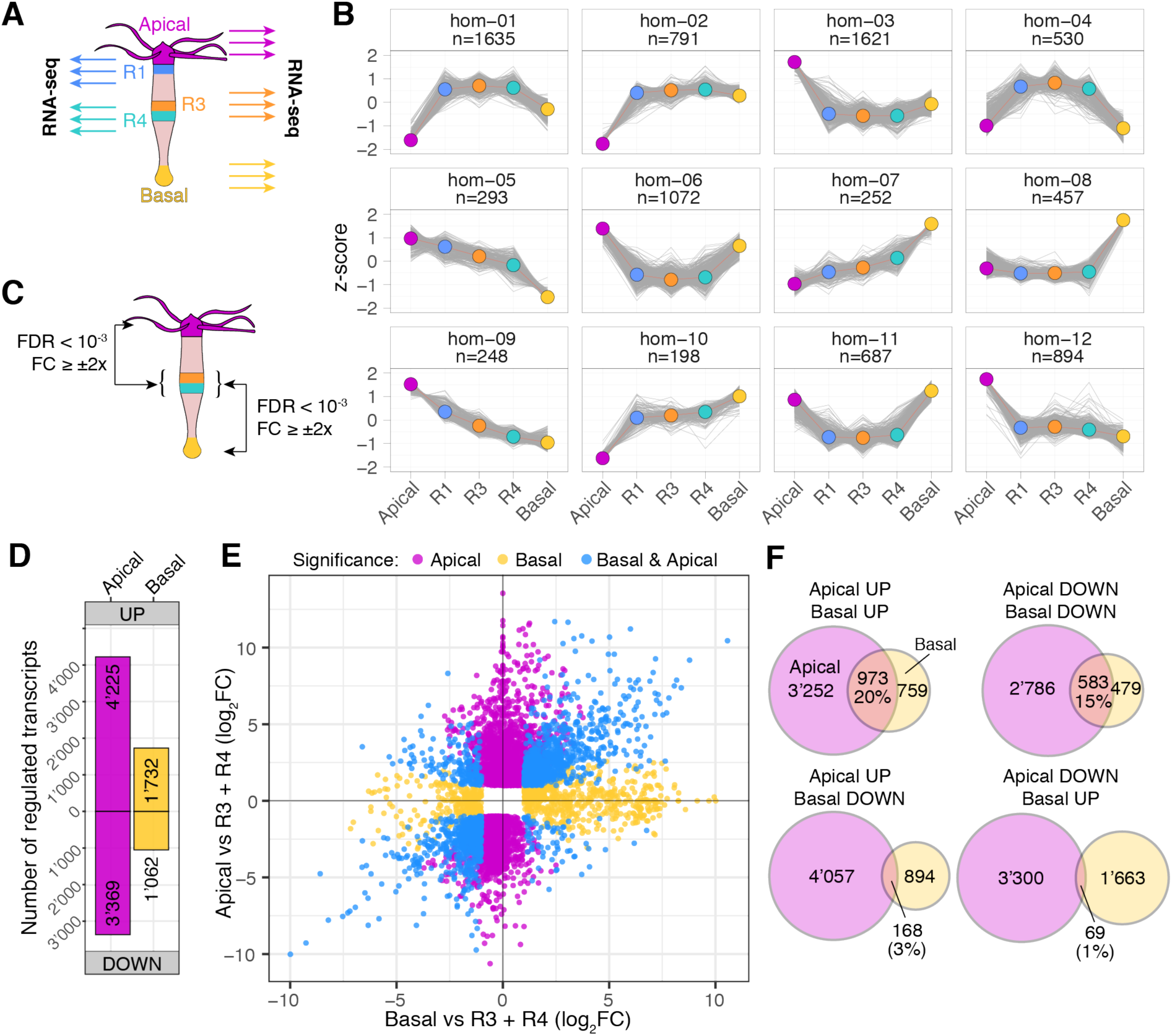
Global transcriptomic analysis of the spatial gene regulations along the *Hydra* body axis. **(A)** Regions sampled for the homeostatic analysis. The slices measure 5% to 10% of animal length (about 500 µm). **(B)**Hierarchical clustering of 8’678 transcripts with variations along the body column (Likelihood ratio test with FDR < 10^−3^ and coefficient of variation ≥ 0.1, see materials and methods for details). **(C)** Schematics of the regions used to compare levels at extremities against the central body column using a Wald test. Only transcripts with False Discovery Rate (FDR) < 10^−3^ and absolute Fold change (FC) ≥ 2 were selected. The triplicates from both R3 and R4 were considered to represent the central body column. **(D)** Transcripts up-or downregulated (FDR < 10^−3^, FC ≥ 2x) in the apical (7’594) or basal (2’794) regions when compared to R3 and R4 regions of the body column. **(E)** Detailed view of FC for transcripts regulated in the apical and basal regions. Colors indicate statistical significances for one or both regions. Transcripts without a significant value in both conditions are omitted from the graph. **(F)** Relationships between transcripts up- and down-regulated in apical or basal regions. Note the larger overlap when regulations occur in the same direction.

To obtain a precise inventory of the genes with differential expression at the extremities, we performed a statistical analysis between apical or basal levels compared to the central body column (Fig. 1C, 1D, **Table S2**). Transcripts were considered as regulated if exhibiting an absolute fold change (FC) superior to 2x and an FDR < 10^−3^ (Fig 1C). We found 7’594 transcripts modulated in the apical region (4’225 up, 3’369 down) but only 2’794 (1’732 up, 1’062 down) in the basal one, indicating a 2.5 times excess in high-apical versus high-basal transcripts (Fig. 1D). Noteworthy, more than 50% of the 1’732 high-basal transcripts are also high-apical (973), indicating that only 759 transcripts are exclusively high-basal while 3’252 are high-apical only, i.e. a 4.3x excess for the latter (Fig. 1E, 1F).

### Genes showing modulated expression at the extremities

Out of 7’594 transcripts with elevated apical levels, 6’911 (91%) sequences match Panther families (Mi et al., 2005), 4’698 (62%) encode Pfam domains (Finn et al., 2014) and 4’709 (62%) show some similarity with human proteins (blastx, E-value < 10^−3^). Overall, 92% (7’014) of these transcripts could be annotated with one or more method (**Table S2**). Some of the high-apical genes detected here were previously identified, as for example *nematocilin* B (<u>seq14014</u>), a cnidarian-specific gene that encodes a component of the cnidocil (Hwang et al., 2008) and shows the highest apical to body column ratio (FC > 10^4^). Out of the 11 known *Wnt* genes identified in *Hydra* (Lengfeld et al., 2009), nine of them are indeed highly expressed apically with FC over the body column values ranging from 5x for *Wnt11* (<u>seq36604</u>) up to 850x for *Wnt16* (<u>seq02507</u>) (Fig. 2A, **Fig. S5**). As anticipated, two *Wnt* genes are not modulated along the body axis, *Wnt2*, known to be expressed only in early stages of budding, and *Wnt9/10b* whose expression was detected only after pharmacological over-activation of the Wnt/β-catenin pathway (Lengfeld et al., 2009).

**Figure 2.**
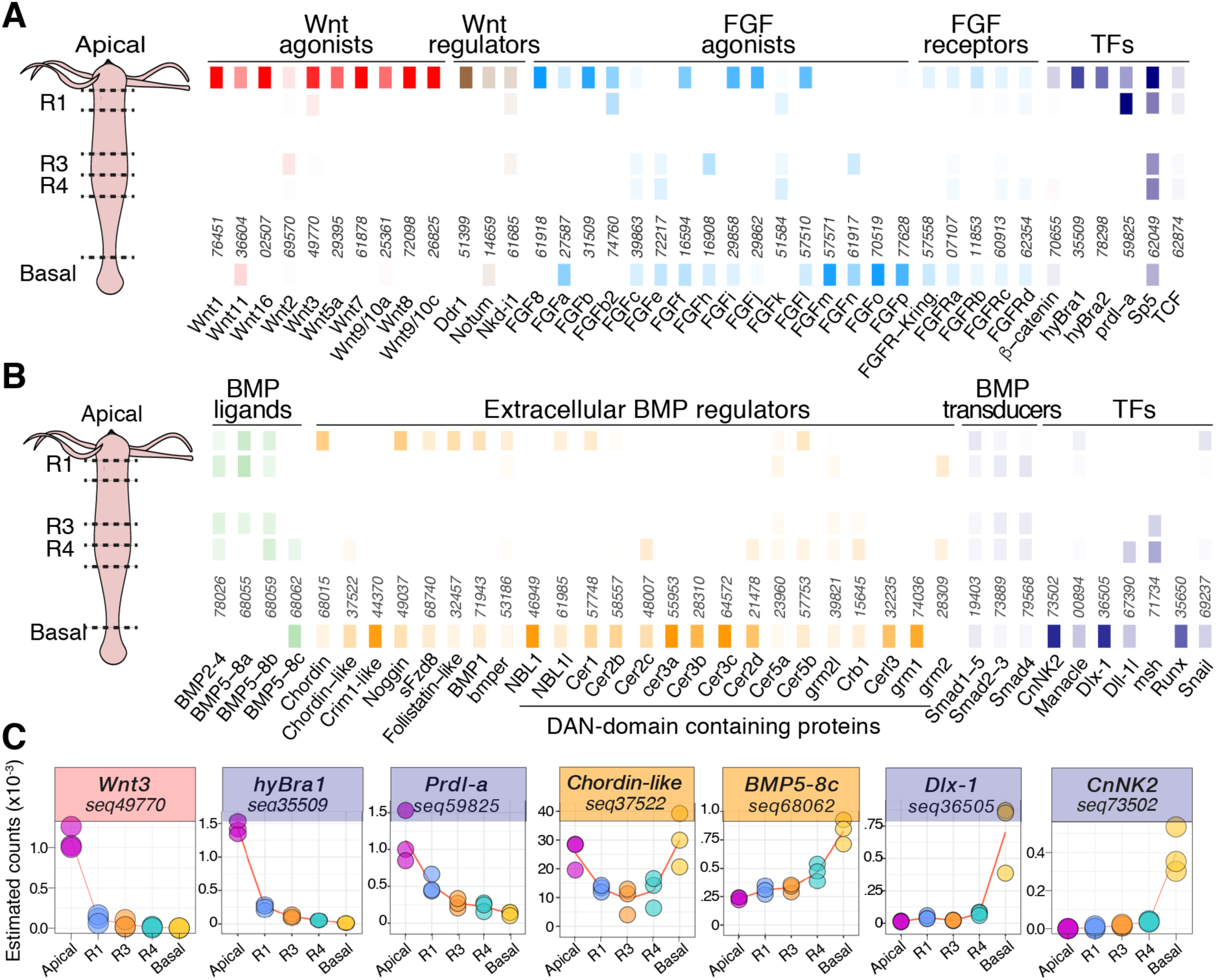
Apical and basal markers from Wnt, FGF and BMP signaling pathways. **(A, B)** Heatmap of spatial expression levels of genes encoding Wnt agonists (red), Wnt signaling regulators (brown), FGF agonists and receptors (light blue), BMP ligands (green), extracellular BMP regulators (orange), Smad proteins (grey) and transcription factors (TFs, purple). The color opacities correlate with relative levels of expression in each of the five sampled regions. Pfam annotation performed on the reference transcriptome identified 34 sequences encoding DAN domains, which were manually curated for redundancy. Three of these sequences do not encode a signal peptide according to phobius (http://phobius.sbc.su.se/) and were removed, yielding a total of 20 sequences. Numbers written vertically indicate ANs on HydrATLAS (type seq-----). **(C)** Spatial expression profiles of selected apical and basal markers.

By coupling the spatial analysis with PFAM domain annotation we identified known as well as previously undescribed genes from transcription factor families, namely T-box/Brachyury (PF00907.21), homeodomain (PF00046.28), Forkhead box (PF00250.17) and Zn finger (PF00096.25) proteins. Among high-apical genes, we identified: the two Brachyury homologs, *HyBra1* (<u>seq35509</u>) and *HyBra2* (<u>seq78298</u>) (Technau and Bode, 1999; Bielen et al., 2007), 14 homeodomain-containing proteins including *cnot* (<u>*seq10145*</u>) (Schummer et al., 1992), *prdl-a* (<u>seq59828</u>) (Gauchat et al. 1998), *HyAlx* (<u>seq42164</u>) (Smith et al., 2000), five Forkhead box proteins, and seven Zn finger proteins (Fig. 2A, **Fig. S5**). Consistently with a study reporting that 13/17 genes encoding Thrombospondin type 1 repeats (TSR, PF00090.18) are predominantly expressed in the hypostome (Hamaguchi-Hamada et al., 2016), we found 14 TSR genes among the high-apical genes (**Table S2**).

Among the low-apical genes (i.e. lower apical levels when compared to the central body column) we also found genes encoding transcription factors: 11 homeoproteins, 12 Forkhead box proteins, eight Zinc-finger proteins (**Table S2**). Regarding the 1’732 transcripts with high expression in the basal region compared to the central body column, eight encode homeodomain-containing proteins including Cn*NK2* (<u>seq73502</u>) (Grens et al. 1996; Thomsen et al. 2004; Siebert et al. 2005), *Dlx* (<u>seq67390</u>), *Dlx1* (<u>seq36505</u>) (Gauchat et al. 2000), one a Forkhead box protein, and eight Zinc-finger proteins. But we also find *sFRP-3* (<u>seq51367</u>) and *Fzd8* (<u>seq68740</u>) (Fig. 2B, 2C, **Fig. S6**), encoding orthologs of the human secreted frizzled-related proteins (Wenger and Galliot, 2013). sFRPs can inhibit Wnt/β-catenin signaling as well as BMP signaling via BMP-1/Tolloid interactions (Lee et al., 2006; Bovolenta et al., 2008).

In total, 973 transcripts were detected as “bipolar”, i.e. statistically significantly up-regulated in both the apical and the basal regions compared to the central body column, sometimes with transcripts levels several times higher at one or the other extremities (see for example *HyAlx* <u>seq42164</u>, **Fig. S5**). *Noggin* (<u>seq49037</u>, Fig. 2B, Fig. 3, **Fig. S6**) that was identified as a BMP inhibitor involved in the Spemann organizer in *Xenopus* (Spemann and Mangold, 1924; Zimmerman et al., 1996) exhibits strongly elevated expression values in both the basal and the apical regions, and virtually no expression in the central body column. The *Hydra* Noggin protein is able to induce a secondary axis when misexpressed in *Xenopus* embryos and is thus considered as homologous to the vertebrate Noggin (Chandramore et al., 2010). Similarly *Chordin-like* (<u>seq37522</u>), which also encodes an inhibitor of BMP signaling (Rentzsch et al., 2007), is expressed at higher levels at the extremities although with a lower significance for the apical region (FDR = 0.0019, Fig. 2B, **Fig. S6**). We also verified the expression pattern of a series of eight genes by whole mount in situ hybridization and indeed we found *Nkd* and *Ddr1* (<u>seq51399</u>) apically expressed, *Noggin* and *Cer1* (<u>seq57748</u>) bipolar, *NBL1* (<u>seq46949</u>), *Crim-1* (<u>seq44370</u>) and *sFzd8* basally expressed (Fig. 3, **Fig. S11**). Overall, these qRNA-seq results are consistent with the results obtained in independent expression studies performed at the single gene level and thus support our experimental design. Therefore, we assume that the proposed pipeline analysis provides reliable transcriptome-wide results.

**Figure 3.**
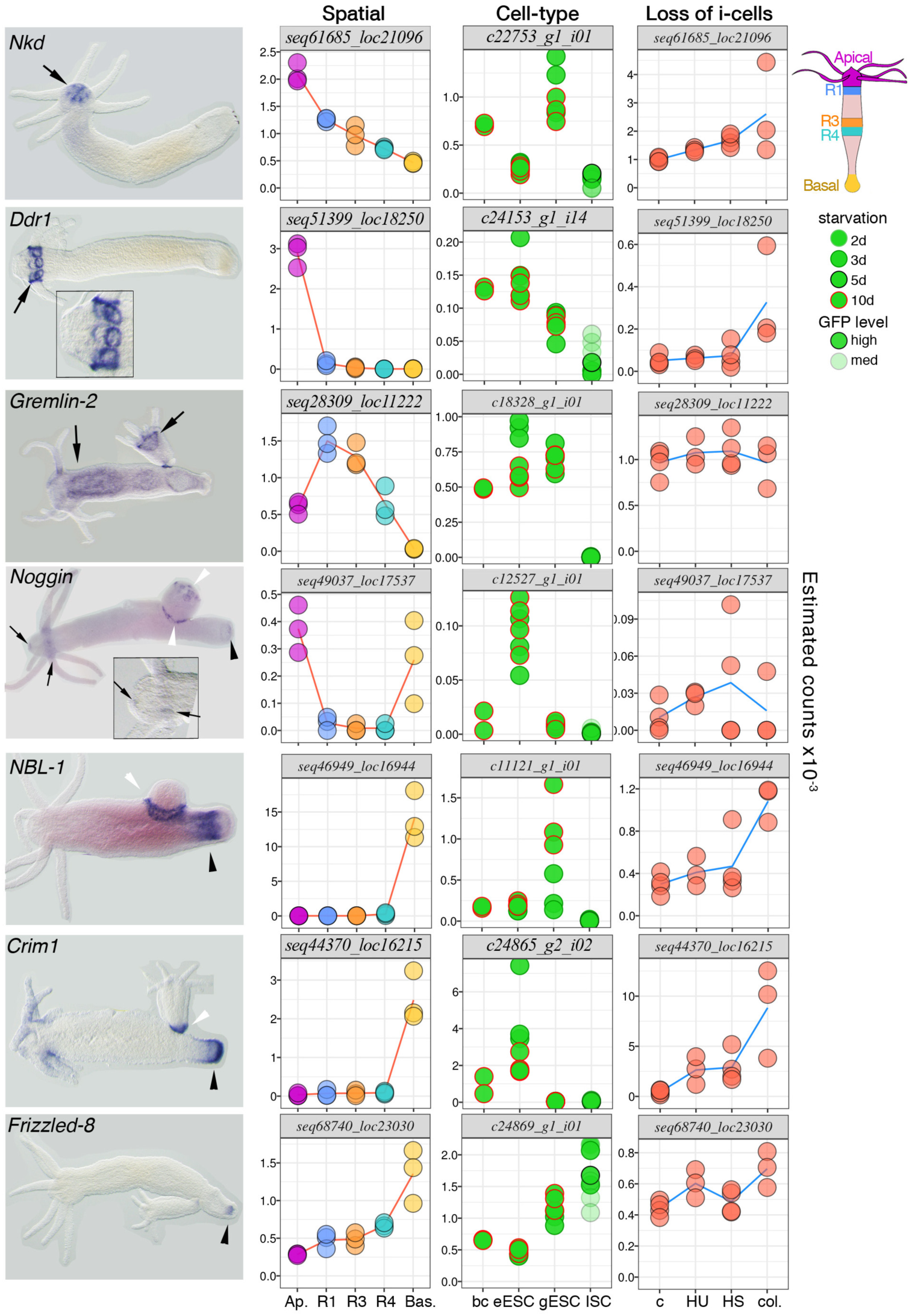
Homeostatic expression patterns of putative regulators of the head and foot organizer. For each gene data are shown on a row, with from left to right the expression pattern, the spatial, cell-type and i-cell loss RNA-seq profiles. Black arrows: apical expression, white arrowheads: expression in the bud or the budding zone, black arrowheads: basal expression. Abbreviations: Ap.: apical, Bas.: basal, c: control untreated animals, eESC: epidermal epithelial stem cells, gESC: gastrodermal epithelial stem cells, ISC: interstitial stem cells. See more animals in supplementary Figure S12.

### Genes with a graded spatial expression patterns during homeostasis

The main unanticipated observation is the high number of transcripts with seemingly graded expression patterns along the animal (793 transcripts), either apical to basal (541 transcripts, clusters hom-05, hom-09) or basal to apical (252 transcripts, hom-07) (**Table-S5**). Although no previous study was specifically designed to capture graded gene expression profiles along *Hydra* axis in an unbiased manner, the expression pattern of some transcripts identified here were already known, such as the Dickkopf homolog *Dkk1/2/4-C* (<u>seq73113</u>) (Augustin et al., 2006) or *BMP2-4* (<u>seq78026</u>) (Watanabe et al., 2014). Looking at individual profiles reveals that the slopes of these gradients along the gastric cavity can either be readily visible (e.g. *BarHl_*<u>seq15014</u>; *TCF* (<u>seq62874</u>), *gremlin2* (<u>seq28309</u>) as confirmed by in situ hybridization – ISH-), or masked when detected by ISH by expression of higher magnitudes at one extremity such as *hyBra1* (<u>seq35509</u>), *prdl-a* (<u>seq59828</u>) or *Wnt3* (<u>seq49770</u>) at the apex, or *NK2* (<u>seq73502</u>) at the basis (Fig. 2, **Fig. S5, S6, Table S1**). Interestingly, the expression of the cell autonomous antagonist of Wnt/β-catenin named *naked cuticle* in *Drosophila* (*Nkd*, <u>seq61685</u>) is graded apical to basal at all five positions (Fig. 3), consistent with a graded regulation of Wnt signaling activity along the *Hydra* body axis as observed in the planarian worm *S. mediterranea* (Stuckemann et al., 2017).

As transcripts showing a typical basal to apical graded distribution and encoding signaling components, we found the ligand *BMP5-8c* (<u>seq68062</u>, Fig. 2C). Three distinct *BMP5-8* genes were identified in *Hydra* (Reinhardt et al. 2004, Watanabe et al., 2014), *BMP5-8b* transcripts being detected at the tentacle level and in the peduncle (Reinhardt et al. 2004). We found *BMP5-8a* (<u>seq68055</u>) gradually expressed apical to basal as shown by Watanabe and coll. but could not confirm the bipolar expression of *BMP5-8b* (<u>seq68059</u>) (Fig. 2B, **Fig. S6**). The sequence alignment identifies three distinct proteins, two of them with two isoforms (**Fig. S7**). The corresponding genes likely arose through tandem duplications as revealed by the phylogenetic and genomic analyses (**Fig. S8, S9**). We also found several potential inhibitors of BMP signaling gradually expressed basal to apical such as *Cer1, Cer2c, Cer3b, Cer3c, Cer3d* (Fig. 2B, **Fig. S6**) and a phylogenetic reconstruction identified 17 distinct genes expressing Cerberus-Gremlin-DAN related proteins in *Hydra* (**Fig. S10**). All together these results suggest that BMP signaling is potentially active at both poles, although with different components and different regulators.

### Cell-type specific gene regulations in homeostatic *Hydra*

Next, we quantified transcripts levels in *Hydra* stem cell populations (Fig. 4A). Each population of stem cells was enriched using fluorescence activated cell sorting (FACS) on tissue from the central body column of transgenic *Hydra* constitutively expressing eGFP either in gastrodermal epithelial cells (gESC, Wittlieb et al., 2006), or in epidermal epithelial cells (eESC, Anton-Erxleben et al., 2009), or in interstitial cells (ISC, Khalturin et al., 2007). These enriched stem cells were then subjected to RNA-seq together with unsorted gastric columns (GC) from intact animals. Using these stem cell specific and GC data, we developed a metric that we call the enrichment score (**Fig. S12A, S12B**, see materials and methods for technical details), which allows to effectively recognize transcripts expressed in interstitial cell derivatives. Briefly, high expression levels of a given transcript in GC samples relative to the levels observed in the three enriched stem cell populations suggest that this transcript was not enriched during the FACS procedure, and thus is predominantly expressed in cell types that are distinct from the stem cell populations.

**Figure 4.**
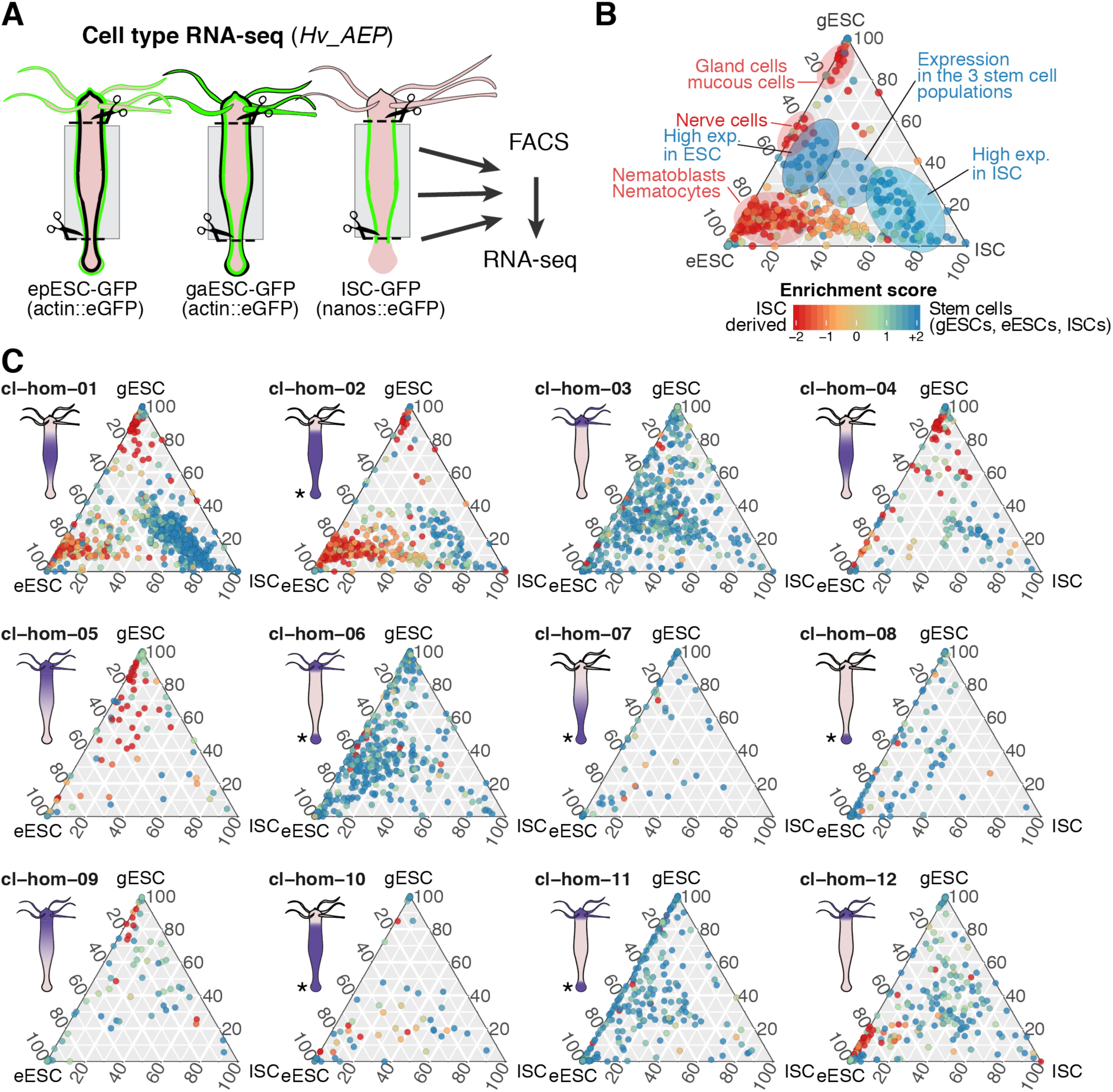
Relationships between spatial and cell-type gene regulations in homeostatic *Hydra*. **(A)** Schematic view of the procedure used to measure gene expression in stem cells constitutively expressing GFP sorted by flow cytometry. This expression was compared to that measured in unsorted gastric columns from intact animals (not depicted). Legend as in Figure 3. **(B)** Ternary plot showing the typical distribution of transcripts from targeted stem cells (blue) or from ISC derivatives (red). **(C)** Ternary plots showing the cell-type distribution of genes defined in the 12 homeostatic clusters shown in Fig. 1B. For each transcript, the enrichment score indicated by the color code was calculated as the log-ratio of the sum of eESC, gESC and ISC values over the value in the unsorted gastric sample. High values (green/blue colors) indicate a strong enrichment compared to full body column tissues in at least one of the factions considered, while low values (orange/red colors) indicate that a transcript is expressed at higher levels in one or many cell types likely not sampled in the eESC, gESC, or ISC fraction. Purple staining on the *Hydra* anatomies are predictions of expression patterns based on the spatial RNA-seq data. *As the lower part of the peduncle was sampled together with the basal disk, it is not possible to discriminate between these two regions using the strategy presented here.

To validate this hypothesis, we used a dataset of transcripts attributed to specific cell types by ISH (Hwang et al., 2007). As we expected, transcripts expressed in gland cells, nematoblasts, nematocytes and nerve cells had negative enrichment scores (**Fig. S12C**). Furthermore, their relative proportions revealed by their positions on ternary plots (**Fig. S12C**) allowed to attribute them to distinct ISCs derivatives. We observed that gland/mucous cells co-segregated with gastrodermal epithelial cells, while nematoblasts, nematocytes segregated with epidermal ones, and nerve cells equally from both, consistent with their tissue layer localizations (**Fig. S12D**). In summary, these data allow to quantify expression levels for transcripts expressed in gESCs, eESCs, ISCs, and to attribute cell type specificity to transcripts expressed in the three stem cell populations as well as in interstitial derivatives in the gastric region (Fig. 4B, **Fig. S12E**, **Table S3**). Finally, we evaluated the presence of ISCs in the epithelial stem cells populations. For this, we assessed the relative expression of *Cnnos1*, which is exclusively expressed in ISCs, in gESCs and eESCs. Our data indicate a moderate presence, with 6% of total *Cnnos1* counts in eESCs and 14% in gESCs (**Fig. S12F**).

### Linking spatial to cell-type specific gene regulations in homeostatic *Hydra*

To investigate the relationship between spatial and cell type expression, we combined transcripts clustered by spatial expression patterns (Fig. 1B) with their cell type specific data (Fig. 4C). We found the vast majority of transcripts predominantly expressed in ISC when expression is restricted to the gastric region (hom-1, and in some extent to hom-04), as anticipated from the absence of ISCs at *Hydra* extremities. However, some ISC transcripts could also be found in hom-02, which can be explained by the sampling of a portion of the peduncle together with the basal disk (Fig. 1A). A gene ontology analysis of transcripts expressed in the stem cells (i.e. with enrichment scores >1) and predominantly expressed in the central body column (hom-01) shows a high enrichment of biological process related to DNA replication, DNA repair, cell cycle (**Table S4**). This result is consistent with the fact that cells in S-phase are largely restricted to the central body column and mostly consist in interstitial stem cells (David and Campbell, 1972; Campbell and David, 1974; Holstein and David, 1990; Buzgariu et al., 2014). The transcripts with low enrichment scores in the gESC fractions, correspond to the known distribution of the gland cells, i.e. restricted to the gastric body column (hom-1/2/4), or to mucous cells, which are present in the upper body column and the hypostome (hom-5/9) (Hwang et al., 2007)(Fig. 4C).

Intrigued by the high numbers of transcripts expressed in a graded manner (Fig. 1A, hom-05/07/09), we had a closer look at their cell-type expression patterns. For this, we first selected transcripts with robust graded expression in the gastric region, in which all triplicates of estimated counts in R1 were higher than those in R3, and R3 triplicate values were higher than those in R4 for apical to basal gradation, and similarly R1 < R3 < R4 estimated counts for basal to apical gradation (**Table S5**). Using these criteria, we found 180 transcripts graded apical to basal and 66 transcripts graded basal to apical. In general, graded expression was present along the full body length of the animal, whereas in some cases gene expression appeared to be regulated independently at the extremities (**Fig. S13**, highlighted boxes). The association with cell-type specific data indicates that such graded transcripts are predominantly detected in gland cells, and to a lesser extent in epithelial cells, either gastrodermal or epidermal. These results indicate that ISCs, at least those where the *Cnnos1* promoter is active, do not express transcripts in a graded manner while gland cells and epithelial cells from both layers do.

To evidence the impact of ISCs and interstitial derivatives on ESCs, we took advantage of the possibility to eliminate the interstitial fast cycling cells with cytotoxic drugs (hydroxyurea – HU-, colchicine) or heat-shock in the thermosensitive *Hv_sf-1* strain to perform a transcriptomic analysis at various time points after such treatments (**Fig. S14**). The kinetics of transcript depletion upon interstitial cell loss shows a rapid effect of the HU treatment, already visible by day-4 when the treatment ends, a loss that gets amplified over the next 7 days. We also noted a limited depletion in the gESC fraction after colchicine treatment, most likely transcripts expressed in the gland cells sampled as contaminant of the gESC fraction (**Fig. S13**). The loss of interstitial cells leads to changes in the gene expression programs of the surviving epithelial cells (Wenger et al., 2016), here visible in several candidate regulators of apical or basal organizers (Fig. 3).

### Global gene expression patterns during basal and apical regeneration

To obtain a detailed view of the modulations in gene expression during regeneration, we performed time course experiments in three different regenerative contexts (Fig. 5A). After an initial amputation at mid-gastric level or below the tentacle ring (corresponding to 50% and 80% of the animal length, respectively), animals were left to regenerate for time periods ranging from 30 minutes to 48 hours (**Fig. S1**) and regenerating tips were sampled. That way, gene expression levels of all *Hydra* transcripts from three regenerative conditions can be systematically followed over time and compared: basal regeneration from the mid-gastric position (BR-50), apical regeneration from the mid-gastric (AR-50) or sub-tentacular (AR-80) positions (see graphical examples of the data generated for *Wnt3* on **Fig. S2**, graphical data for all transcripts are available on the <u>Hydratlas</u> server).

**Figure 5.**
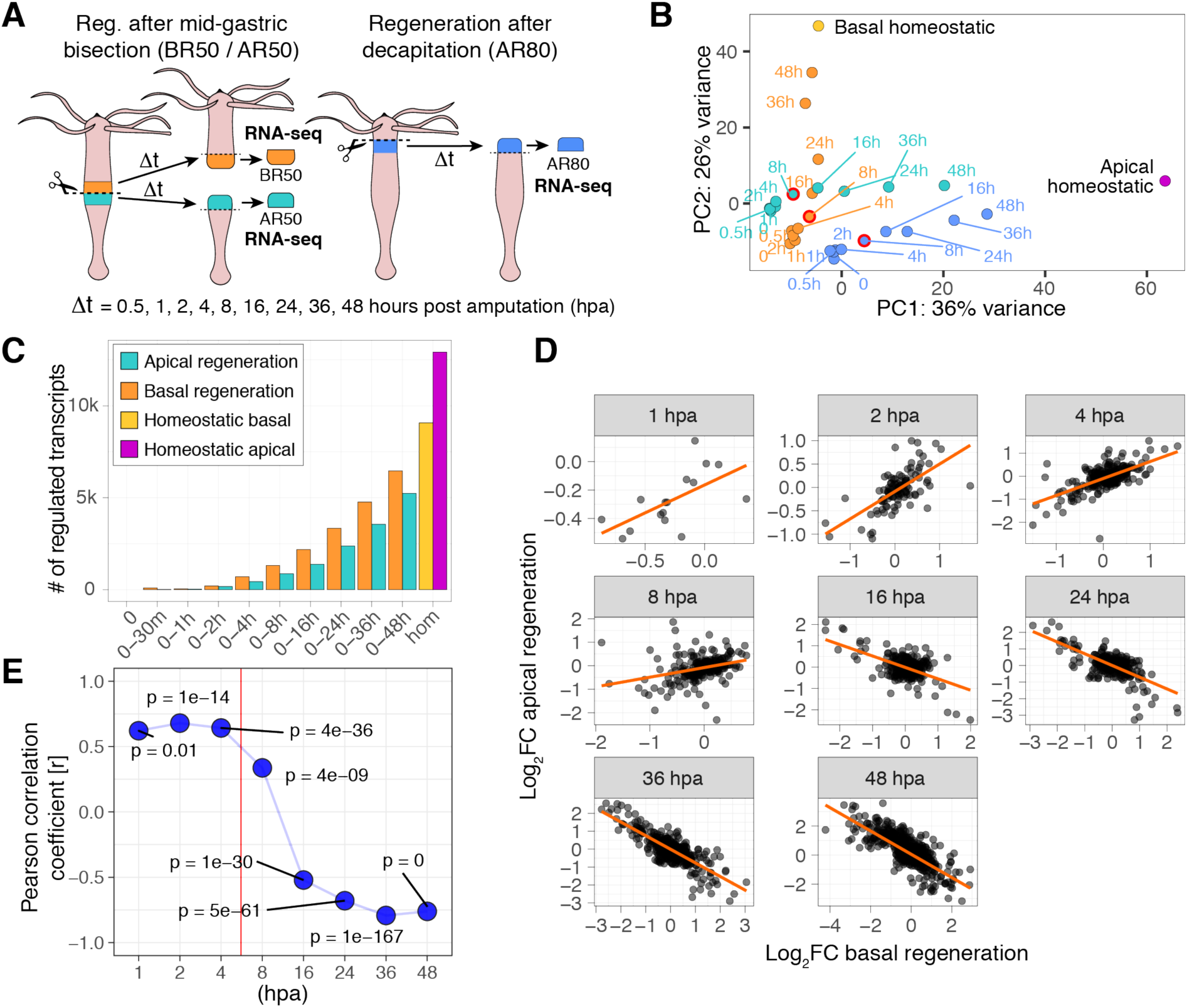
Global transcriptomic analysis of the temporal gene regulations identified during *Hydra* regeneration. **(A)** Scheme depicting the experimental procedure to identify modulations in gene expression in regenerating tips at 9 distinct time-points of apical (AR) and basal regeneration (BR), either after mid-gastric bisection (AR50, BR50) or after decapitation (AR80). Gene expression levels measured in the upper (R1) and central body column regions (R3 or R4) dissected in the absence of any regeneration were used as time 0 (t0) conditions for AR80, BR50, and AR50. All 30 conditions were sampled in three independent biological replicates, resulting in 90 samples. **(B)** PCA analysis of samples collected after mid-gastric bisection (green), decapitation (blue) and in basal-regenerating tips (orange). First movements towards the structure to be regenerated are detected by 8 hpa in all three regenerative conditions (red-circled dots). At 48 hpa, values of basal-regenerating tips almost reach the value of the basal homeostatic condition, which is not the case when comparing the respective positions of the apical regenerating tips at 48 hpa and the homeostatic apex. **(C)** Timings of gene regulations for transcripts with predominant apical (7’594) or basal (2’794) expression shown in Fig. 1D-1F. Each bar represents the results of one of the 20 likelihood-ratio tests performed using the samples included in the intervals ranging from t0 to the indicated time, FDR < 10^−3^. **(D)** Correspondences between significant apical and basal FCs during the progression of regeneration (FCs calculated over t0 values). Each dot represents one transcript and red lines fitted linear models. Only transcripts with statistically significant FC in both conditions are represented and used for the modeling. **(E)** Summary plot of the Pearson correlations calculated from data represented in panel D showing an onset of change in gene regulation between AR50 and BR50 occurring between 4 and 8 hpa, followed by a massive uncoupling between and 8 and 16 hpa. P-values specifying the significance of the measured Pearson correlations are indicated (p).

We first performed a global assessment of the sample behaviors by principal component analysis (PCA) using transcripts with regulations detected by ANOVA and with a variation over time of more than 10% of the average expression (2’998 transcripts, Fig. 5B, **Fig. S4**). This analysis provides several observations: firstly, all three cases considered here (AR-50, AR-80 and BR-50) have different origins, i.e. initial conditions are different indicating a level of heterogeneity in gene expression along the body column. Secondly, for any of the three considered conditions, samples for regeneration time-points between 0 and 4 hours post-amputation (hpa) are overlapping, and then start diverge towards the final endpoints by 8 hpa (Fig. 5B, circled-red dots). Thus, a detectable change in gene expression among the 2’998 transcripts considered here occurs between 4 and 8 hpa, a time interval where regenerating tips start acquiring gene expressions more related to the final structures that will be regenerated, and which also corresponds to the initial rise in head organizer capacity (MacWilliams, 1983b). Thirdly, the trajectories of the samples over time are almost completely orthogonal between apical and basal regeneration, indicating that, among the selected genes, only a limited amount of gene modulations are shared between apical and basal regeneration processes. Fourthly, the AR-50 and AR-80 regeneration trajectories are roughly parallel and then tend to converge after 48 hpa. Fifthly, after 48 hours of regeneration, a time point initially chosen because the morphological structures are already in place for both apical and basal regeneration, the PCA analysis indicate that the basal regeneration condition almost reached the full “homeostatic” signature, whereas more time is required for apical regeneration.

Previous reports in *Hydra* (Wenger et al., 2014; Petersen et al., 2015; Gufler et al., 2018) and planaria (Wenemoser et al., 2012) identified genes activated early after injury, irrespective of the position and expected fate of the stump. These genes were associated with wound-healing, immunity, or even patterning as β-catenin targets that were shown to behave similarly at the transcriptional level during the first hours of regeneration (Gufler et al., 2018). To establish if these injury-induced modulations are global or restricted to a subset of transcripts, we investigated the mutual relationship between regulations occurring during basal and apical regeneration in a transcriptome-wide manner. We retrieved transcripts with statistically significant fold changes between t0 and sampling time points ranging from 1 hpa to 48 hpa for both apical and basal regeneration (Fig. 5C). We then measured for each time point the direction and strength of relationships between fold changes over time in both regenerative conditions.

The slopes of the regression lines are positive at early time points and evolves towards negative values with longer regeneration times (orange line, Fig. 5D) while the correlations between apical and basal fold changes follows a similar trend (Fig. 5D, **Fig. S15B**). Therefore, the fold changes of transcripts regulated in both apical and basal regeneration conditions are highly positively correlated up to 4 hpa, whereas the strength of the correlation starts to decrease between 4 and 8 hpa, and this relationship is dramatically reversed by 16 hpa, maintained negative at all subsequent time-points (Fig. 5E). In other terms, for genes regulated in both AR-50 and BR-50 conditions, gene modulations observed up to 4 hpa, and most gene modulations occurring up to 8 hpa occur in a context-independent manner.

To confirm the observation that a large number of genes involved in regeneration are regulated after 48 hpa, we identified all transcripts significantly modulated over time intervals ranging from t0 to 30 min, or from t0 up to complete regeneration (Fig. 5C). Reasoning that fully regenerated apical and basal regions are undistinguishable from homeostatic apical and basal regions, we considered that homeostatic conditions for these regions represent conditions when regeneration is complete. Accordingly, we found 12’919 and 9’075 transcripts modulated during apical and basal regeneration respectively; however, between t0 and 48 hpa, only 40% (5’240) are significantly modulated during AR-50, versus 71% (6’466) during BR-50.

### Comparison of apical and basal regeneration identifies generic (impulses) and context dependent (state-transitions) modulations in gene expression

Typical transcriptional responses to environmental stimuli include impulse and sustained (“state-transitions”) patterns (Yosef and Regev, 2011). In regenerating *Hydra*, we found both types of patterns regulated either positively or negatively, with variable onset timings (Fig. 6, **Fig. S16**). An impulse-type response during early regeneration was previously described as generic, i.e. similar between AR-50 and BR-50, for 43 transcripts. These transcripts were annotated as immunity-related (Wenger et al., 2014), including components of the p38/MAPK pathway such as *Jun* and *Fos*, but also encoding transposable element domains (Petersen et al., 2015). Here, we used an impulse-based model (Fischer et al., 2017) to classify gene expression patterns obtained during *Hydra* regeneration. Using stringent parameters (FDR < 10^−5^), we detected 683 and 744 impulse-regulated transcripts during AR-50 and/or BR-50 respectively (Fig. 6A, 6C, **Table S6**). Transcripts detected as sustained with onsets starting during the first 48 hours of the regeneration process amounted to 3’231 for AR-50 and 3’826 for BR-50 (Fig. 6B, 6E, **Table S6**).

**Figure 6.**
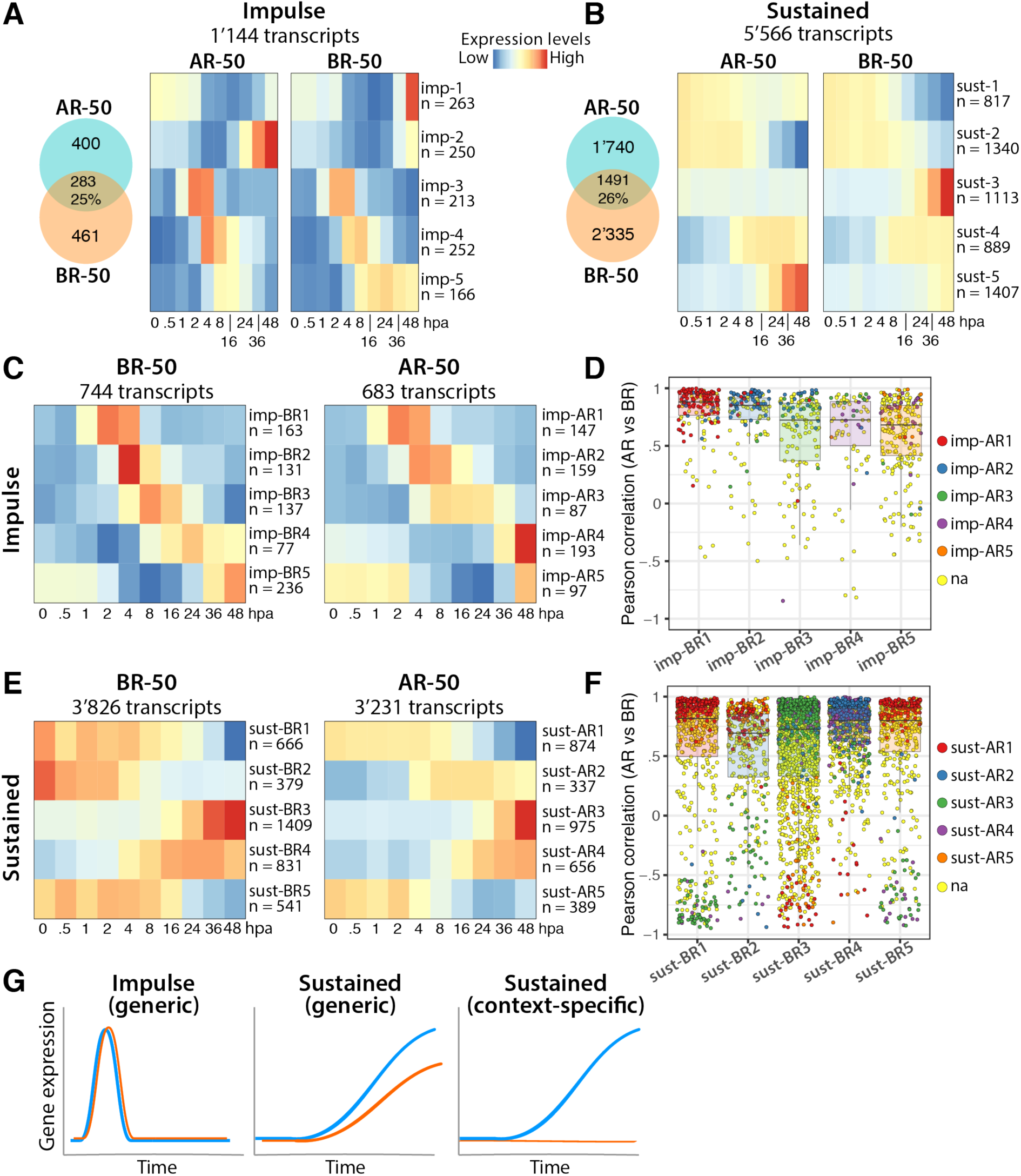
Impulse and sustained expression patterns observed during *Hydra* regeneration. **(A, B)** Number of genes showing impulse (imp, **A**) or sustained (sust, **B**) patterns during AR-50 and/or BR-50. **(C, E)** Heatmaps representing the evolution over time of gene expression classified as impulse **(C)** or sustained **(E)** in AR-50 and BR-50 conditions. (**D, F**) Scatterplot representations of the pairwise Pearson correlations between the gene expression levels of transcripts assigned to impulse-type (D) or sustained-type (F) regulations. Transcripts selected as impulse or sustained were clustered by the k-means method applied on standardized read counts and median values per clusters are shown. The x-axis indicates the BR-50 clusters taken as references, the colors represent the AR-50 cluster identity and each dot represent a Pearson correlation coefficient calculated for each transcript between the AR and BR conditions. na: not attributed as impulse or sustained patterns in AR50. For sustained transcripts, a majority of patterns are specific to AR-50 or BR-50 (clusters 1, 2, 5) although few exhibit a similar behavior (clusters 3, 4). **(G)** Summary scheme: impulse regulations occur at early timepoints and are shared between AR-50 (blue) and BR-50 (orange), left panel. Most sustained regulations occur at later timepoints and can be either generic (central panel), or context-specific (right panel).

To understand the relationships that occur between the behaviors of regulated transcripts during AR-50 and BR-50, we clustered together the AR-50 and BR-50 patterns from the impulse-type transcripts on the one hand (1’144 transcripts) and the sustained-type transcripts (5’566 transcripts) on the other hand (Fig. 6A, 6B). We observe that transcripts classified as impulses tend to overall behave similarly in the two regenerative contexts, while some sustained clusters behave very differently, in particular clusters sust-5 and sust-3 that contain late up-regulated transcripts. The earliest group of regulated transcripts is imp-3, with an upregulation visible by one hpa, peaking at 2-4 hpa (Fig. 6A), whereas the earliest onset of sustained regulation is seen at 4 hpa (sust-4, Fig. 6B). Overall, these observations, which are in agreement with those presented in the previous section (Fig. 5C, 5D), clearly establish that early regulations are predominantly generic impulses, while later regulations are dominated by sustained behaviors that can be generic or context-specific (Fig. 6G).

Next, we clustered independently the patterns detected as impulse and sustained in the AR-50 and BR-50 conditions (Fig. 6D-F, **Fig. S17**). Among clusters classified as impulse, we observed similar timings of regulation and correlations of gene expressions patterns over time in BR-50 and AR-50 (Fig. 6C, 6D). For example, transcripts from the cluster imp-BR1 that exhibit the earliest global regulation during basal regeneration, are also regulated during AR-50, found in the imp-AR1 cluster that exhibits the earliest modulation (Fig. 6D, red dots). The high levels of co-variations (high Pearson correlation coefficients) for the transcripts categorized as impulse indicate that a majority of these transcripts behave similarly during apical and basal regeneration. However, a limited number of transcripts were detected as impulse-regulated in only one of the two conditions examined (Fig. 6D, yellow circles in the column imp-BR1). We manually inspected these patterns and found that they fall into two categories: either an impulse visually comparable in both conditions but statistically significant in a single one according to ImpulseDE2, or transcripts can show an early impulse profile and at a later time point a striking sustained profile, thus classified as sustained rather than impulse. As a general rule for impulse-type patterns, the earliest is the onset of regulation, the more likely it is that the gene expression pattern is generic.

Concerning the sustained up-regulations, two clusters, sust-BR3 and sust-BR4, are detected in the BR-50 condition with onsets at 16 and 4 hpa respectively (Fig. 6E). Looking at the sust-BR4 transcripts, we found 337 also regulated during AR-50, affiliated to the clusters sust-AR2 and sust-AR4 with 172 and 165 transcripts respectively (Fig. 6F, blue and purple circles) and 16 transcripts attributed to sust-AR1 and sust-AR5, i.e. clusters characterized by a negative regulation during apical regeneration. Regarding the 1’409 sust-BR3 transcripts, we found 932 (66%) not regulated in the AR-50 condition, 401 detected in the sust-AR3 and sust-AR4 clusters, and 58 (4%) in the negatively regulated sust-AR1 and sust-AR5 clusters (Fig. 6E-6F). These results imply that a series of genes can be concomitantly submitted to opposite regulations in the head- and foot-regenerating tips.

### Early genes with sustained expression are candidate effectors of the transition from gastric tissue to apical and basal organizing centers

To identify candidate genes involved in the setting up and the maintenance of the apical and basal organizers, we searched for genes with early and sustained expression levels when compared to expression levels in the initial tissue (i.e. region R3 for BR and region R4 for AR, Wald test, FC ± 2x, FDR < 10^−3^) at every time point from 8 hpa to full regeneration, the latter being assumed to be represented by levels in extremities of intact animals (Fig. 7A). Only a few genes display an onset of sustained regeneration (OSR) before 4 hpa while the number of genes initiating a sustained regulation increases over time, except for up-regulated transcripts in the BR-50 condition that become less numerous at 48 hpa.

**Figure 7.**
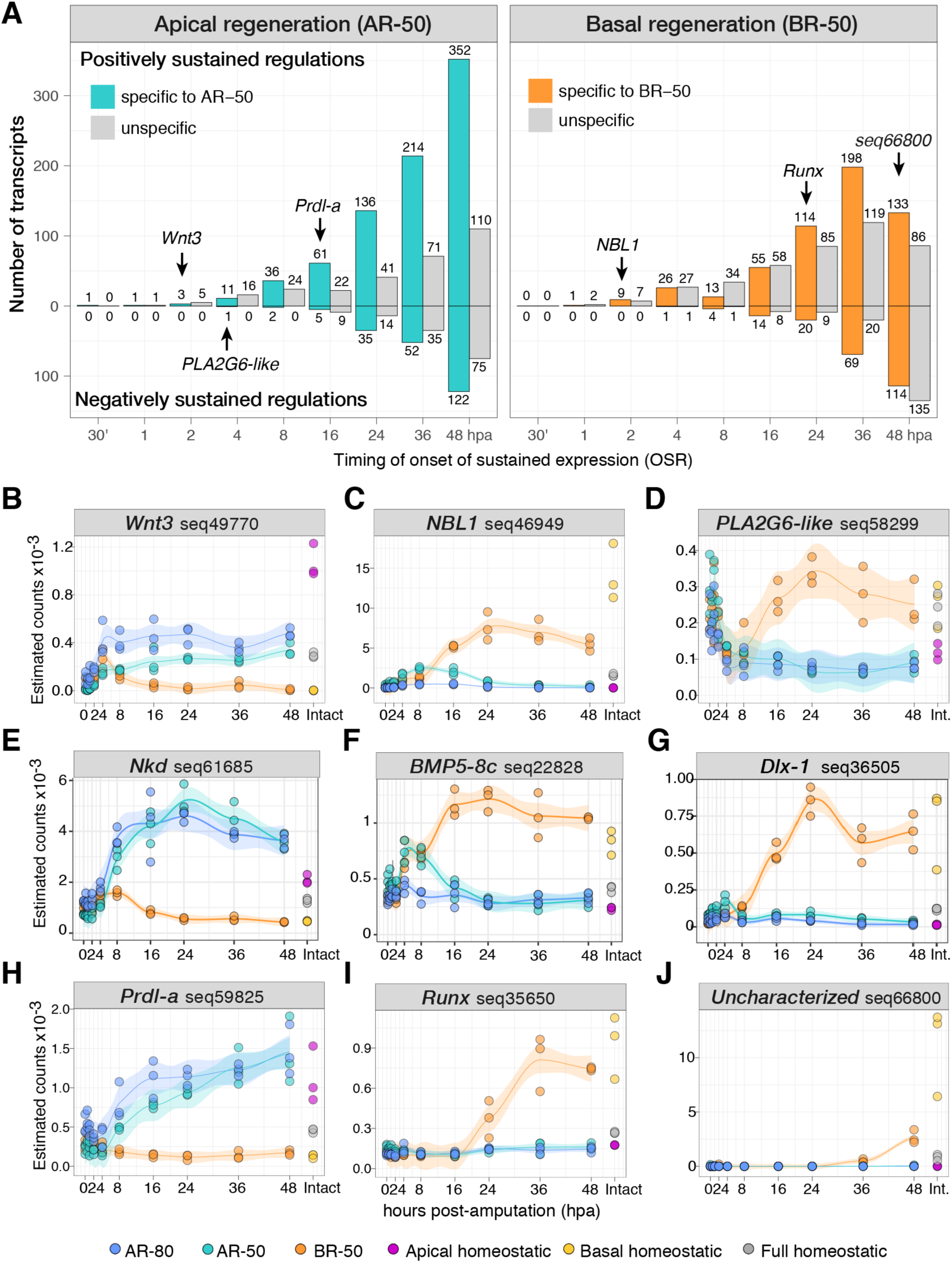
Stepwise increase in the number of sustained regulations during apical and basal regeneration. **(A)** The onset of a sustained regulation (OSR) is defined as the first timepoint when a regulation (positive or negative) is recorded with a minimal two-fold change (FC > ±2x, FDR < 10^−3^) compared to the time-0 value and maintained qualitatively similar at all subsequent timepoints and in the final structure to be regenerated. The time-0 value was measured in regions R4 for AR-50 and R3 for BR-50, dissected in the absence of regeneration (Fig. 1A, 2B, 2C). Numbers in the graph indicate for each time point the number of genes that initiate a sustained regulation, grey bars correspond to counts of transcripts showing an unspecific sustained regulation, i.e. detected in the same direction (positive or negative) in both AR-50 and BR-50 conditions, color bars represent transcripts with distinct sustained regulations in AR-50 and BR-50, either in a sustained manner in only one condition or with opposite ones. **(B-J)** Examples of regeneration profiles. **(B, C)** *Wnt3* and *NBL1* are detected as sustained up-regulated at 2 hpa for AR-50 and BR-50, respectively. **(D)** *PLA2G6-like* is down-regulated by 4 hpa in the AR-50 condition and transiently negative but not sustained in the BR-50 condition. **(E, H)** In AR-50, OSR is detected at 8 hpa for *Nkd* and 16 hpa for *prdl-a*. **(F, G, I)** In BR-50, OSR is detected at 16 hpa for *BMP5-8c* and *Dlx-1*, 24 hpa for *Runx*. **(J)** A late uncharacterized basal transcript with OSR detected at 48 hpa. Green: AR-50, blue: AR-80, orange: BR-50, purple: apical, yellow: basal, grey: full animal.

For apical regeneration, we identified 133 transcripts (106 loci) satisfying these criteria, comprising only six that were consistently down-regulated (**Fig. S19**, **Table S7**). Among the three genes already up-regulated at 2 hpa, we found *Wnt3* (Fig. 7A, 7B, Fig. 8A), which is central in setting up the head organizer (Broun et al., 2005; Lengfeld et al., 2009; Nakamura et al., 2011). The remaining two contigs, which contain predicted peptide signals, correspond to proteins uncharacterized in *Hydra*. One of them harbors an insulin growth factor binding protein domain (IGFBP) and an antistasin domain, whereas a spliced isoform of the same gene lacks the IGFBP domain. Although the equivalent of the mammalian insulin could not be detected in *Hydra* tissues, *Hydra* reacts to insulin treatment through a reaction mediated by an insulin-like receptor (Steele et al., 1996). Several insulin-like peptides have been identified and proposed to mimic the role of insulin in higher organisms (Fujisawa and Hayakawa, 2012). Furthermore, insulin and insulin-like growth factors are important modulators of the RAS/MAPK pathway in bilaterians (Weng et al., 2001; Fujisawa and Hayakawa, 2012).

**Figure 8.**
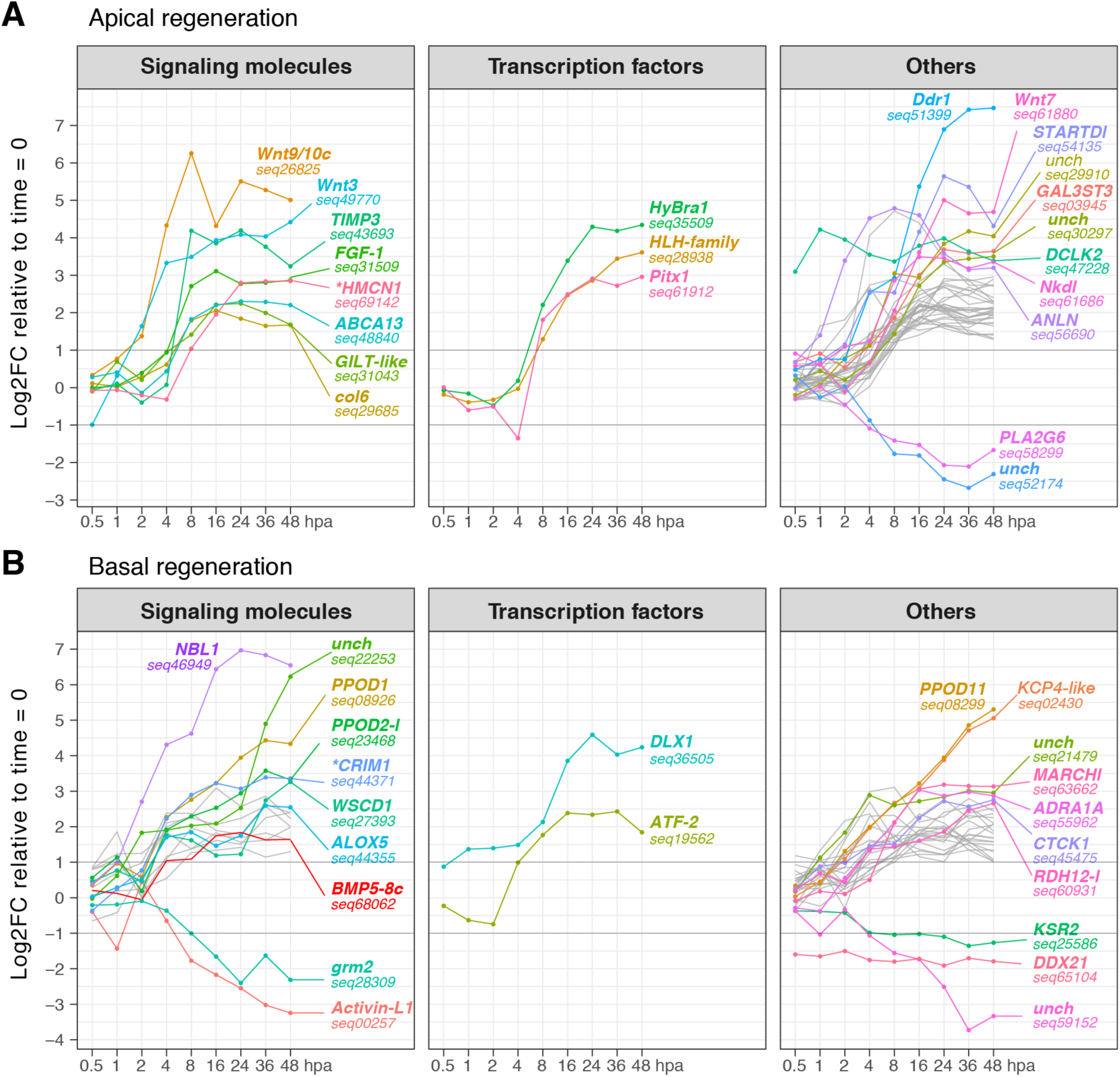
Early sustained gene regulations identify potential candidates of apical and basal organizers. Genes showing an OSR before 8 hpa as defined in Fig. 7 (FC > ± 2x, FDR < 10^−3^) and the highest FC during apical **(A)** or basal **(B)** regeneration between 8 and 48 hpa. Gene names were first found on UniProt after blastx or blastp on the 11’133 *Hydra* protein sequences available (E < 10^−50^), then similarly searched on NCBI if not found on UniProt, otherwise deduced from the closest human sequence as indicated by asterisks * (E < 10^−3^). If no related characterized sequence could be found, the gene was named uncharacterized (Unch.). *BMP5-8c* (seq68062) is indicated as a red line.

Concerning the early-sustained basal genes that candidate for patterning (**Fig. S20**), we found *NBL1*, which encodes a unique DAN domain (Fig. 7A, 7C, Fig. 8B). The *Hydra* NBL1 sequence shows the highest sequence similarity with the NBL1 protein family but does not branch as a true NBL1 ortholog in phylogenetic trees (**Fig. S10**). In mammals as in *Hydra*, the DAN superfamily also includes the Cerberus and Gremlin gene families (Katoh and Katoh, 2006). Proteins harboring DAN domains are known to interact with BMP signaling (Kattamuri et al., 2012; Nolan et al., 2013), and are also common players of Wnt and Nodal signaling (Katoh and Katoh, 2006; Watanabe et al., 2014).

Among the early-sustained genes encoding secreted proteins, we found *FGF-1* in head-regenerating tips, potentially contributing to the activity of the head organizer (Fig. 8A), and *BMP5-8c* in foot-regenerating tips, potentially responsible for setting up the basal organizer through interactions with *NBL1* (Fig. 8B). The number of transcription factor genes showing a two-fold sustained early-upregulation is limited: *hyBra1, Pitx1* and a *HLH* member for apical regeneration, *Dlx-1* and *ATF-2* for basal regeneration (Fig. 8, Fig. 7D). Among the other regulators, we identified in AR-50 *Nkd*, a potential intra-cellular inhibitor of Wnt/β-catenin signaling as mentioned above (Fig. 7E, Fig. 8A). We also analyzed the profiles of the early expressed genes with bipolar distribution, i.e. activated in both the AR-50 and the BR-50 condition. We found a single transcript activated by four hours *Ets1* (<u>seq73274</u>), which is predominantly expressed in the basal region in homeostatic condition. The transcription factor HyEts1 belongs to the ERG/FLI1 families of Ets containing proteins, which, in vertebrates, are target of the ERK/MAPK pathway and cooperate with members of the AP-1 complex to activate Ras-responsive elements (Wasylyk et al., 1998; Yordy and Muise-Helmericks, 2000; Foulds et al., 2004; Hollenhorst, 2012). Assessing Ets1 activity might lead to some valuable insights on the asymmetric activation of the MAPK pathway after mid-gastric bisection (Galliot, 2013).

### BMP signaling as a potential driver of the basal organizing center

The basal region of the *Hydra* polyp acts as an organizer center, inducing an ectopic foot when grafted onto the body column of a host (Fig. 9A). As *BMP5-8c* is predominantly expressed in the basal region and the only growth factor gene defined as early-sustained, we postulated that BMP5-8c might act as a positive component of the basal organizer also named “foot activator”. All components of BMP signaling are expressed in *Hydra* and the phosphorylation sites required for the nuclear translocation of the transcriptional co-activator Smad are conserved (Fig. 9B, 9C). In fact, we detected higher levels of the phosphorylated form of Smad (p-Smad) in lower halves of intact animals, implying a higher BMP activity in the basal region of the animal (Fig. 9D). Among the putative BMP inhibitors, *NBL1* shows a rapid up-regulation after bisection, only sustained in foot-regenerating tips (Fig. 7C, 9E). In homeostatic conditions, *NBL1* is expressed at maximal levels in the peduncle showing a sharp boundary with the basal disc but a graded basal to apical extension towards the body column (Fig. 3, **Fig. S11**). All these results fulfill the criteria of a basal organizer relying on the interactions between positive BMP signaling and NBL1 inhibition.

**Figure 9.**
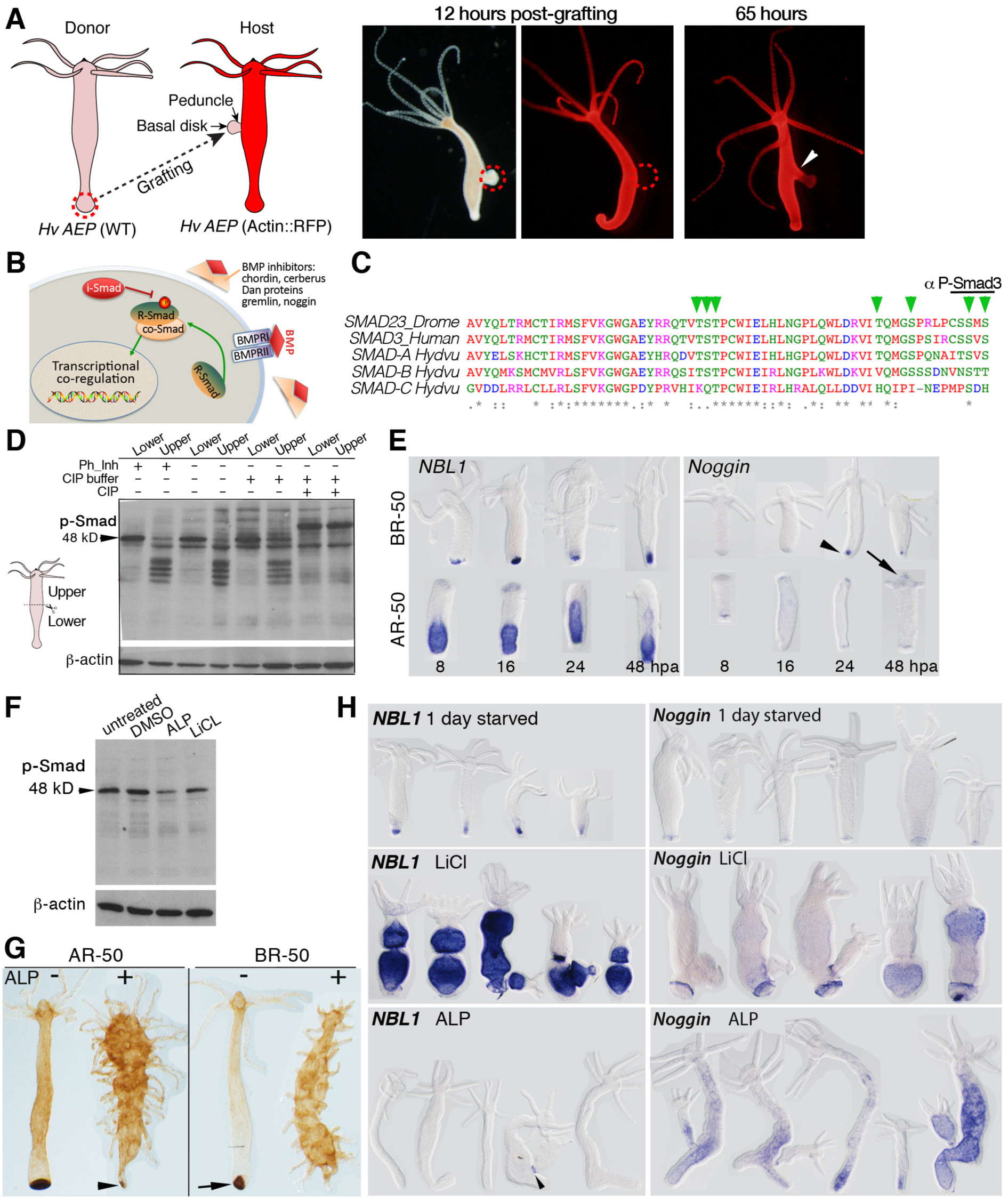
The *Hydra* basal organizer and BMP signaling in the basal region. **(A)** Activity of the basal organizer as evidenced by a transplantation experiment from the basal region of a non-transgenic *Hy_AEP* animal onto the body column of a transgenic *Hy_AEP* animal constitutively expressing RFP. Note the ectopic growth of a basal region that includes cells from the host (arrowhead). **(B)** Schematic view of the activation and inhibition of the BMP signaling pathway. Extra-cellular BMP inhibitors identified in bilaterians are expressed in *Hydra*. **(C)** Partial alignment of the *Drosophila*, human and *Hydra* Smad sequences showing the Ser/Thr phosphorylation sites involved in Smad activation (green arrowheads). **(D)** Western Blot analysis of phosho-Smad (p-Smad) levels in extracts from upper and lower halves of intact *Hydra*. The 48 kDa band that corresponds to the predicted *Hydra* Smad size (red arrow) disappears upon phosphatase (CIP) treatment. A higher p-Smad level in lower halves suggests that BMP signaling is more active there. **(E)** *NBL1* and *Noggin* expression in the halves regenerating apically (AR-50) and basally (BR-50) after mid-gastric bisection. Note the sustained *NBL1* expression in foot-regenerating tips, the peak of *Noggin* expression in the foot-regenerating tips at 24 hpa (arrowheads) and the sustained *Noggin* expression at the apex (arrow). **(F)** Western Blot analysis of p-Smad levels in intact animals exposed either for two days to Alsterpaullone (ALP) or for one week to LiCl. **(G)** Peroxidase activity is a marker of the basal disc in *Hydra*. Peroxidase activity is measured here in animals exposed or not to ALP for two days, amputated mid-gastric and left to regenerate for 4 days. ALP-treated animals differentiate tentacles all along their body axis. Note the disappearance of the constitutive basal activity in animals regenerating their head (AR-50, arrowhead) and the lack of basal differentiation in the upper half undergoing basal regeneration (BR-50). **(H)** *NBL1* and *Noggin* expression patterns in intact animals exposed to LiCl for four days or to ALP for two days. Note the expansion of the *NBL1* domain all along the body column after LiCl exposure and the suppression of *NBL1* expression after ALP except at the base of a mature bud (arrowhead). By contrast, *Noggin* expression expands basically upon LiCl treatment and in the central body column upon ALP.

As the activity of the basal organizer is supposed to be repressed by the vicinity of the head organizer, we used two pharmacological agents, LiCl known to induce ectopic foot formation when given continuously over several days (Hassel and Berking, 1990; Jantzen et al. 1998) and Alsterpaullone (ALP) that constitutively activates Wnt/β-catenin signaling along the body column (Broun et al. 2005). These two distinct conditions are expected to differently impact on BMP signaling, LiCl expanding the basal organizer activity along the body column, and ALP repressing it by inducing ectopic apical organizers along the body column. We found that both drugs decrease the level of pSmad (Fig. 9E), which is surprising for LiCL but fully expected after a two-day ALP treatment. To better visualize the effect of ALP on the basal organizer, we used the peroxidase activity as a marker of basal differentiation (Hoffmeister and Schaller, 1985) (Fig. 9F, **Fig. S21**). Animals were bisected after a two-day ALP treatment and the peroxidase activity was detected four days later. All ALP-treated animals showed a large number of ectopic tentacles along the body column as expected, but no peroxidase activity at the basal pole, indicating that basal differentiation did not take place during basal regeneration (Fig. 9F, right panel, **Fig. S21B, D**). In lower halves undergoing apical regeneration, we noticed that the strong peroxidase staining of the basal disc was lost in 50% of the animals, suggesting a progressive dedifferentiation of the basal region (Fig. 9F, arrowhead, **Fig. S21B**). In terms of gene regulations, we found the *NBL1* expression domain dramatically expanded along the whole-body column immediately after a seven-day LiCl treatment and completely abolished after a two-day ALP exposure (Fig. 9G). When we analyzed the drug-induced regulation of the bipolar gene *Noggin*, we detected an up-regulation at the boundary of the basal disc upon LiCl exposure and a widespread up-regulation along the body column in response to ALP, thus a quite different response than that displayed by *NBL1*. All together these results support the idea that *NBL1* regulation reflects well the cross-talk between the apical and the basal organizers, a high level of *NBL1* expression being locally required for maintenance of the basal disc and for foot formation.

## DISCUSSION

### Strengths and limitations of the RNA-seq approach

This study demonstrates that the quantitative RNA-seq approach, validated by a series of previously published expression patterns, is well suited to establish the precise mapping of the genetic regulations in organisms that maintain a highly dynamic homeostasis, and to monitor in time and space the changes in gene expression that accompany complex developmental processes. The preparation of identical samples in three independent experiments performed at distinct periods of the year provides robust and highly significant RNA-seq results. Indeed, the observed variations between sample values are limited although more frequently observed in the R4 region, which corresponds to the budding zone of the animal where the budding potential fluctuates and the expression of genes involved in budding highly dynamic (Boettger and Hassel, 2012). Therefore, we consider that most of the observed variations between the samples do have a physiological significance. Also, the RNA-seq quantification highlights one of the drawbacks of the ISH method. Due to the large range of RNA-seq sensitivity, we could detect low, but consistent expression of genes in locations where they were thought to be completely silent. The best example is the graded expression of *Wnt3* along the *Hydra* body column, with expression levels ~100x lower in the body column when compared to the apical region.

Concerning the homeostatic patterns, it should be noted that the apical and basal regions used in this study are not strictly restricted to the extremities and include some cells from the upper body column (immediately below the tentacle ring) for the apical samples, and some cells from the lowest part for the peduncle for the foot samples. The cell-type analyses performed here on the three stem cell populations sorted by flow cytometry do not have the resolution of single-cell sequencing as recently made available in *Hydra* (Siebert et al., 2018). Nevertheless, the quantitative comparison with intact tissue and the complementary analyses on tissues depleted of their interstitial cells allow deductions that are highly informative on gene expression in interstitial derivatives such as gland cells, nerve cells and nematoblasts/nematocytes. One limitation of this study concerns the analysis of the temporal regulations during three distinct types of regenerative contexts (AR-80, AR-50, BR-50) as RNAs were extracted from limited pieces of tissues, i.e. the regenerating tips that were compared to tissues taken at the same position along the body column from non-regenerating animals. As a consequence, potential systemic responses such as seen in planarians (Wenemoser and Reddien, 2010) and zebrafish (Lepilina et al., 2006) could not be distinguished here from the local responses restricted to the regenerating tip.

### Wound-healing in *Hydra* relies on evolutionarily-conserved processes

Restoring complex tissue structures upon injury involves the reactivation of developmental processes that are specific to the structure to be regenerated. This ability is quite mosaic across the animal kingdom (Bely and Nyberg, 2010). In contrast, stress and wound repair responses are ubiquitous in multicellular organisms, but have also been shown to be remarkably context-independent (Gurtner et al., 2008). Here we identified 1’144 transcripts with impulse regulation, and among them, two context-independent early waves peaking at 2-4 hpa (imp-03, imp-04) and 4-8 hpa, likely involved in the wound healing process as deduced from the fact that they are consistently upregulated whatever the regenerative program. These two waves include genes involved (or annotated) in the immune response as well as genes involved in the remodeling of the extra-cellular matrix. These results indicate that in *Hydra* as in bilaterians wound healing occurs independently of the type of regeneration program that is activated at the same time.

### Confirmation of Wnt3 as the key component of the head organizer

Lateral transplantation experiments have evidenced the presence of a head organizer in intact animals as well as the *de novo* formation of a head organizer in head-regenerating tips within the 12 hours following bisection (Browne, 1909; Yao, 1945; Webster and Wolpert, 1966; MacWilliams, 1983b; Takano and Sugiyama, 1983), reviewed in (Shimizu, 2012; Vogg et al., 2016). Similarly, transplantation experiments evidenced the presence of a foot organizer active at the basal pole (Hicklin and Wolpert, 1973). Classically, the activity of organizers relies on the release of substances named morphogens, which act differently on the recipient cells according to their level of expression (Gurdon and Bourillot, 2001; Plouhinec et al., 2013; Shimozono et al., 2013). In zebrafish embryos, such gradients appear essential to control the formation of embryonic axes (Xu et al., 2014). As graded biological activities were directly quantified in *Hydra* by lateral transplantation experiments (Berking, 1979; MacWilliams, 1983b, Shimizu, 2012), it is expected that cells sense signals distributed in a graded fashion, and respond to these positional signals by altering their gene expression program.

The observed graded expression of *Wnt3* together with the earliest up-regulation after mid-gastric bisection fully supports the expected roles of Wnt3 as the key activator component of the head organizer (Hobmayer et al. 2000; Broun et al., 2005; Guder et al., 2006; Lengfeld et al. 2009; Nakamura et al., 2011). The graded expression of *Wnt3* uncovered in this study implies different levels of activity along the body column with high levels maintaining the head organizer active at the apex of the animal in homeostatic conditions, and lower levels supporting *Wnt3* auto-activation through β-catenin signaling at any level along the body column. The fact that among the 3’231 genes that show a sustained up-regulation during head regeneration, *Wnt3* appears as the earliest one, is a striking observation that strongly supports its role in the re-establishment of an active molecular head organizer at the apical regenerating tip (Nakamura et al., 2011). In that respect it is surprising to note that the levels of *Wnt3* expression at 48 hpa remain about twice lower than that observed in intact heads, suggesting that the head organizer can take distinct statutes according the surrounding tissues, either homeostatic then involved into head maintenance, or linked to head development.

### Candidate head inhibitors as Wnt3 / β-catenin signaling inhibitors

*Hydra* transplantation experiments also demonstrated that organizers are actually made of two antagonistic components with parallel graded activity along the body axis, one positive named activator and another negative named inhibitor that restricts the activity of the activator (Rand et al. 1926; Webster and Wolpert, 1966; Berking, 1979; MacWilliams, 1983a; MacWilliams, 1983b; Takano and Sugiyama, 1983). The results of extensive transplantation experiments, especially those performed by Webster and Wolpert, provided the basis of a Turing type reaction-diffusion model where the activator and the inhibitor, both diffusible extra-cellular substances, “react auto- and cross-catalytically” (Gierer and Meinhardt, 1972). The short-range activator auto-activates and activates the inhibitor while the long-range inhibitor restricts the activity of the activator.

The recent characterization of the inhibitory component of the head organizer, the transcription factor Sp5, which fulfills a series of criteria, (i) to be a target gene of Wnt/β-catenin signaling, (ii) to be expressed gradually along the apical to basal body axis, (iii) to be reactivated in a sustained manner during head but not foot regeneration, (iv) to inhibit Wnt/β-catenin signaling and (v) to prevent head formation (Vogg et al., 2019) (Fig. 10). Homeostatic *Sp5* expression show three levels, maximal in the apical region, high along the body column but not graded, and low in the basal region, while *Sp5* up-regulation after bisection is only sustained in head-but not foot-regenerating tips. Therefore, the characterization of Sp5 largely relied on the RNA-seq results obtained here. This study also identifies the gene *Naked cuticle* (*Nkd)* as an additional inhibitory candidate of the head organizer, as in mammals and *Drosophila*, Nkd behaves as an intra-cellular negative regulator of β-catenin signaling (Yan et al., 2001), while in *Hydra*, *Nkd* transcripts distribute gradually from the apex and are up-regulated early after bisection in a sustained fashion. In agreement with such intra-cellular scenario, *Wnt3, β-catenin*, *Sp5* and *Nkd* are predominantly expressed in the gastrodermal epithelial cells, which are known to carry the morphogenetic information (Fujisawa, 2003; Shimizu, 2012). Finally, the elegant distribution of *Ddr1 (Discoidin domain receptor 1)* transcripts at the tentacle roots, suggests for this receptor tyrosine kinase interacting with collagens a role in cell migration towards the tentacles, possibly through Wnt5a as identified in mammals (Jonsson and Andersson, 2001).

**Figure 10.**
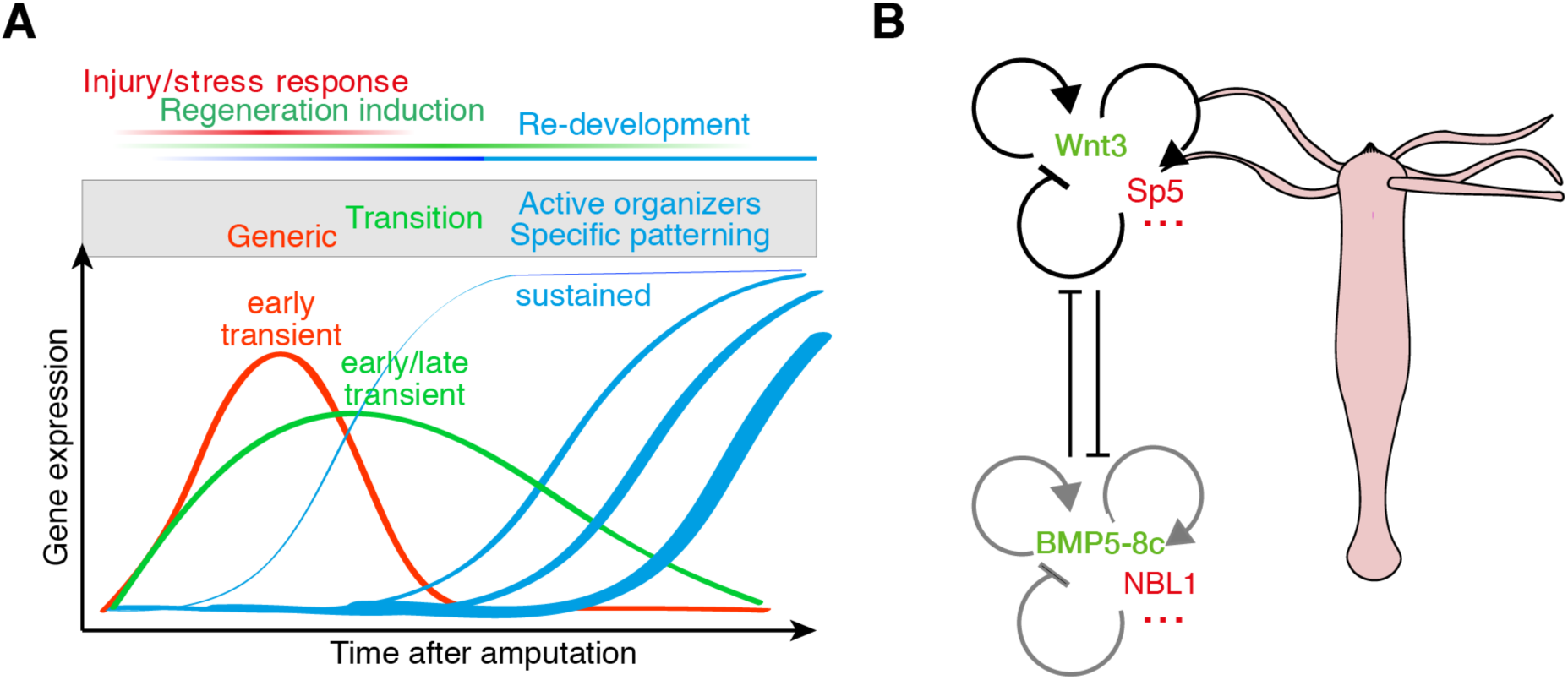
Schemes summing up the gene regulations linked to regeneration and to the head and foot organizers in *Hydra*. **(A)** Summary view of the generic and developmental context-dependent types of gene regulations identified by comparing apical and basal regeneration with a systematic transcriptomic approach. **(B)** Schematic view of the head and foot organizers. Beside the confirmation of the positive and negative actors of the head organizer previously identified, namely Wnt3 and Sp5, this study also identifies potential additional regulators such as Nkd. Concerning the components of the basal organizer, BMP5-8c fulfills the criteria of a positive regulator (basal activator) while NBL1 and additional DAN-domain containing proteins candidate to act as negative regulators of the basal organizer. Grey circular arrows indicate putative regulations.

### BMP signaling as a key regulator of the foot/basal organizer

The combination of spatial-homeostatic and temporal-regenerative transcriptomics allowed us to propose some candidate components for the basal organizer, BMP5-8c as foot/basal activator, and NBL1 as foot/basal inhibitor (Fig. 10). Two related epitheliopeptides, the 49 amino-acid long Hym-346 and the 21 amino-acid long C-term product pedibin, were previously shown to promote foot formation in *Hydra* (Grens et al., 1999). This analysis confirms their predominant expression in gESCs but could trace neither their spatial-homeostatic nor their temporal-regenerative regulation as the number of reads was too low. In *Nematostella*, BMP signaling plays a major role in symmetry breaking during embryogenesis, and a double negative feedback loop between BMPs and chordin could be evidenced, similar to the regulation at work in the dorso-ventral (DV) axis of developing vertebrates (Saina et al. 2009). These results imply that the cross-talk between BMPs and their inhibitors already plays a developmental role in cnidarians, which remains to be dissected at each pole of the *Hydra* polyp and in regenerating animals, especially regarding the cross-talk with β-catenin signaling.

### Relationships between the *Hydra* and vertebrate body axes

The evolutionary relationship between the oral-aboral axis of cnidarians and the antero-posterior (AP) and/or dorso-ventral (DV) axes of vertebrates is unclear, possibly not existing for the *Hydra* axis, rather restricted to the apical pole (Galliot and Miller, 2000). An alternate hypothesis would be that the *Hydra* head organizer is related to the vertebrate AP patterning system, and the foot organizer to the DV system, i.e. including the Spemann organizer in the dorsal side (Rentzsch et al., 2007). In vertebrates, BMP and Wnt signaling levels are integrated to produce the DV (elevated BMP signaling in the ventral region) and the AP (elevated Wnt signaling in the posterior regions) axes (Niehrs, 2004; De Robertis, 2008; Attisano and Wrana, 2013). An interesting proposal was made that the mechanisms involved in dorsoventral patterning of most deuterostomes define the oral-aboral axis of the sea urchin (Duboc et al., 2004), suggesting that the sea urchin case represents a secondary loss of the coupling between the orthogonal AP and DV patterning systems. The results obtained in this study indeed suggest an important role for BMP signaling in the foot/basal organizer in addition to the known roles of Wnt/β-catenin signaling in the head/apical organizer (Fig. 10). The negative cross-talk we identify here between Wnt/β-catenin and BMP signaling suggest that the interactions between these two signaling systems that deliver mutually exclusive positional information predate Cnidaria divergence. The origin of this cross-talk might be either unique and evolutionarily-conserved, or submitted to parallel evolution (Wenger and Galliot, 2013).

## MATERIALS & METHODS

### Spatial RNA-seq during homeostasis and regeneration (*H. vulgaris Jussy*)

*Hydra vulgaris (Hv)* from four distinct strains, Jussy (*Hv_Jussy*), sf-1 (*Hv_sf1*) or AEP (*Hv_AEP*) were used in this study. For *Hv_Jussy* the mass culture was obtained from a single animal collected in Jussy, Canton of Geneva, Switzerland during the summer 2012 (geographic coordinates: 46°15'08.8" N, 6°16'53.5" E). Animals were cultured as described in (Gauchat et al., 2004), at a controlled temperature of 19°C, fed Monday, Wednesday, and Friday mornings with freshly hatched *Artemia* and washed 9 hours after feeding and the following day. All initial tissue bisections started on Monday mornings, before feeding. Each of the following conditions were obtained from 25 *Hydra*: full intact animals, heads and feet of *Hydra* in homeostatic conditions, regenerating tips after either bisection at the 50% or 80% levels (head regeneration), or 50% level (foot regeneration). For the AR-50 condition, 25 animals were bisected at the 50% level in 25 ml Hydra medium (HM), halves corresponding to the BR-50 condition were sorted and placed in another cell-culture dish. For the AR-80 condition, animals were cut below the base of the tentacles and isolated apical regions were discarded. Dissection of the regenerating tips (~250 µm) were performed with scalpels at 0, 30 minutes, 1, 2, 4, 8, 16, 24, 36 and 48 hours after the original bisection.

### Stem cell specific RNA-seq using FACS (*H. vulgaris AEP*)

We used three transgenic lines kindly provided by T. Bosch (Kiel), each constitutively expressing GFP in one of the stem cell populations, and performed fluorescence activated cell sorting (FACS) on dissociated tissues from central body columns, as previously described (Hemmrich et al., 2012; Buzgariu et al., 2014; Wenger et al., 2016). Briefly, we used *H. vulgaris AEP (Hv_AEP)* animals expressing eGFP in their gastrodermal or epidermal epithelial stem cells (gESCs, eESCs) under the control of *Hydra actin* promoter, named Endo-GFP and Ecto-GFP strains respectively (Wittlieb et al., 2006; Anton-Erxleben et al., 2009) or in their interstitial stem cells (ISCs) under the control of the *nanos* promoter, named *Cnnos1*-GFP (Hemmrich et al., 2012). The design of this experiment is comparable to the one presented in Hemmrich et al. (2012), however, the sensitivity of the approach was considerably extended by sequencing and mapping 400 Mio reads using three to four replicates for each sample, compared to a total of about 1.5 Mio without replicates in the previous study. As control values, we used tissues from whole (unsorted) gastric columns (GC) to allow the identification of contaminants in FACS sorted datasets.

### Reference transcriptome preparation and quantification

#### RNA-seq library preparation and high-throughput sequencing

All tissue samples were placed in RNALater (Qiagen) immediately after bisection and total RNAs were extracted within hours with the RNAeasy mini kit (Qiagen). Each condition was sampled in at least three independent biological replicates (except for *Hv_AEP* unsorted gastric data that were sampled in duplicates), representing a total of 99 samples. Input material was 700 ng of total RNA for each sample. Libraries were prepared with the Low Sample TruSeq total RNA preparation protocol from Illumina (San Diego, CA, US) using 15 PCR cycles. Library concentrations were measured with a Q-bit (Life Technologies, Carlsbad, CA, US). Size distribution of fragments was estimated with a 2200 TapeStation (Agilent, Santa Clara, CA, US) and fragments of 350 bp in average were obtained. Pools of 4 or 5 multiplexed libraries were loaded per lane of a HiSeq 2000 sequencer (Illumina) at 8 pM and single-end sequenced using the 100 bp protocol.

#### Reference transcriptomes

The assemblies of the reference transcriptomes used here have been described previously (Wenger et al., 2016). For quantifications, reads from Hv_Sf1(drug exposure RNA-seq) and Hv_Jussy (homeostatic and regeneration RNA-seqs) were mapped to the *Hv Jussy* transcriptome, and reads from *Hv AEP* (cell type specific RNA-seq) to the *Hv AEP* transcriptome.

#### Quantification of reads

Reads processed with cutadapt 1.11 (Martin, 2011) to remove Illumina adapters and known trans-spliced leader (Stover and Steele, 2001; Derelle et al., 2010), were aligned to their respective reference transcriptome (see above) using Salmon 0.7.2 (Patro et al., 2017) and rounded numbers of reads were extracted (“NumReads” columns in quant.sf files provided by Salmon). Library normalization and statistical analyses were done using DESeq2 (Love et al., 2014) and most graphs were produced using the ggplot2 package (Wickham, 2009).

### Sequence annotation in HydrATLAS

Sequences were annotated using the Pfam 31.0 (Finn et al., 2014) and Panther 13 (Mi et al., 2005) databases based on their longest ORF. The occurrence of signal peptide cleavage sites was assessed using SignalP 4.1 (Petersen et al., 2011) and Phobius 1.01. For blast-based annotation, *Hydra* protein and the human reference proteome set were downloaded from UniProt (Dec 2017, 7’076 and 71’775 proteins for *Hydra* and human, respectively). Blastx was used with an E-value threshold of 10^−50^ for Hydra in order to identify identical proteins, or 10^−3^ in human in order to identify related protein families. Annotations for *Hv_Jussy* and *Hv_AEP* are available on figshare.

### Exploratory data analyses

Unless otherwise stated, all analyses were carried out on all transcripts comprising at least one sample of interest with more than 100 reads (**Fig. S4A**).

#### Clustering of homeostatic expression patterns

To detect transcripts with significant variations along the body column, we applied the likelihood ratio test as implemented in DESeq2 (Love et al., 2014) with false discovery rate (FDR) < 10^−3^ and a supplementary threshold to retain only transcripts with variations larger than 10% between the maximum and the minimum median values recorded in all conditions for each transcript (**Fig. S4**). Read counts were variance stabilized using a regularized log transformation method (rlog), and median values per condition and per transcript were computed. To focus on pattern rather than magnitudes, data were standardized (i.e. z-scores were computed so that the average of the read counts equals zero and the standard deviation 1 for each transcripts), and finally clustered using squared Euclidean Distance. Note that clustering applying a squared Euclidean distance on z-scores is strictly equivalent to applying a method based on the Pearson coefficient on the variance stabilized dataset (Berthold and Höppner, 2016).

#### Cell-type specific RNA-seq and ternary plots

On the presented ternary plots (Hamilton, 2017), the enrichment score is calculated as the sum of the expression estimate of the three sorted fractions compared to the unsorted fraction (Fig. 4). That way, transcripts that are present at low levels in one or more of the three samples but are present at high levels in the gastric column are associated with a low enrichment score. Such transcripts originate from cells that were not targeted for sampling but that likely adhered to the eGFP-sorted cells or were present from disrupted cells. To establish their origin, we relied on a curated dataset of specific cell-type transcripts (Hwang et al., 2007). The analysis indicates that contaminant transcripts predominantly sampled in the gESC fraction come from gland cells, those found in the eESC fraction come from nematoblasts and nematocytes while transcripts from nerve cells are found both in the gESC and eESC fractions (Fig. 4B, **Fig. S12**). The results are consistent with the tissue layers where these cells are located. Gene enrichment analysis consists in assessing the frequencies of a particular sequence attribute in a set of interest, to the frequency of the same attribute found in a given background, in this case, the *Hydra* transcriptome. The background used in this study comprises all sequences annotated using PANTHER v13 as described above. In addition, it contains all the identifiers of sequences that were not assigned to a particular family.

### Phylogenetic analyses

Protein sequences were deduced with Translate (http://web.expasy.org/translate/), related sequences were identified with BLASTp on Uniprot/TrEMBL and NCBI databases. Retrieved sequences were then aligned with MUSCLE (http://www.ebi.ac.uk/Tools/msa/muscle/) (Edgar, NAR 2004) or MAFFT (http://mafft.cbrc.jp/alignment/server/), and phylogenetic trees were built with PhyML 3.0 (Guindon et al. 2010).

### HydrATLAS

Detailed results of the RNA-seq data presented in this study are available at https://hydrATLAS.unige.ch

### Data availability

Reference transcriptomes for *Hydra vulgaris Jussy* (<u>GGKF00000000</u>) and *Hydra vulgaris AEP* (<u>GGKH00000000</u>) were deposited on the NCBI nucleotide archive. All Raw Illumina reads and quantified gene expression estimates have been deposited to the Gene Expression Omnibus data repository (GEO) under umbrella accession <u>GSE111534</u>. The 153 raw Illumina read files used in this study have been deposited on the Sequence Read Archive (SRA) with accession numbers ranging from SRX3771898 to SRX3772055, also available through the GEO repository <u>GSE111534</u>. Datasets are available on figshare https://figshare.com/account/home and the code used to produce the figure on GitHub https://github.com/yvanw

Figshare: https://figshare.com/articles/Hv_Jussy_annotation_blast_pfam_panther_/6512957

### Antibodies and Western analyses

WCE were prepared from 20 to 100 polyps starved for two days, washed in HM, resuspended in lysis buffer (SDS 2%, Tris-HCl pH 8.0 100 mM containing a mix of protease (Roche complete cocktail) and phosphatase inhibitors (Biotools cocktail or lab-made cocktail: 8 mM NaF, 20 mM β-glycerophosphate, 10 mM Na3VO4, 0.1 mg/mL PMSF) if not otherwise indicated and passed through a 0.6 × 30 mm syringe needle. For each condition 10-30 μg WCE were loaded on a 12% Rotiphorese^®^ Gel 30 (Roth), transferred to Hybond-P membrane (GE Healthcare) and detected with a rabbit anti-pSMAD antibody (1:1000, ab52903). Extracts were treated with the calf intestinal phosphatase (CIP) as indicated by the supplier (Promega).

### Drug treatments and whole mount in situ hybridization (ISH)

Animals were starved one day before starting the treatment. LiCl 0.5 mM was added to Hydra Medium continuously for 7 days. The animals were fed as usual (3 times a week) and starved for the last two days of treatment. Alsterpaullone (ALP, 5 µM) was given for 2 days to one day starved animals maintained unfed. Whole mount ISH experiments were performed as described in (Wenger et al. 2016).

## Supporting information

HydrATLAS_Supplemental_Data

## Acknowledgements

We are grateful to Marie-Laure Curchod for excellent technical assistance, Mylène Docquier (iGe3 Genomics Platform: ige3.genomics.unige.ch) for excellent support with high-throughput sequencing, Jose Manuel De Abreu Nunes (iGe3 BioSC platform: biosc.unige.ch) and Sylvain Sardy (CAMAS: www.unige.ch/math/en/camas) for helpful discussions. Computations were performed on the Baobab cluster (plone.unige.ch/distic/pub/hpc/baobab_en) at the University of Geneva. This work was supported by the Swiss National Science Foundation (SNF grants 31003A-130337, 31003A_149630, 31003_169930), the NCCR Frontiers in Genetics (Stem Cells & Regeneration work package), the Human Frontier Science Program (HFSP RHP0016-2010), the Claraz donation and the Canton of Geneva.

