## Supplementary material for "Generic and context-dependent gene modulations during *Hydra* whole body regeneration": HydrATLAS_Supplemental_Data

#### Supplementary tables

**Table S1.** Annotations of the sequences corresponding to clustered genes in homeostatic conditions

http://129.194.56.90/STable_Supp_clustering_homeostatic_data_2018-04-20.xlsx

**Table S2.** Annotations of the sequences with regulations at *Hydra*'s extremities compared to the central gastric region (R3 & R4)

http://129.194.56.90/STable_Supp_modulated_in_extremities_2018-04-20.xlsx

**Table S3.** Cross-analysis between the spatial and cell type specific RNA-seqs

http://129.194.56.90/STable_cell_type_attribution_with_clusters_2018-05-17.xlsx

**Table S4.** Gene ontology analysis of stem cells corresponding to clustered genes in homeostatic conditions

http://129.194.56.90/STable_GO_homeostatic_topgo_BP_2018-04-20.xlsx

**Table S5.** Annotations of the sequences strongly graded in the *Hydra* gastric region

http://129.194.56.90/STable_analysis_of_graded_transcripts_2018-05-22.xlsx

**Table S6.** Classification of transcripts into impulse and sustained patterns by ImpulseDE2

http://129.194.56.90/STable_ImpulseDE2_results_2018-05-23.xlsx

**Table S7.** Detection of statistically significant regulations compared to initial condition during AR-50 and BR-50

http://129.194.56.90/STable_significance_of_modulations_by_timepoint_12062018.xlsx

### Supplementary Figures

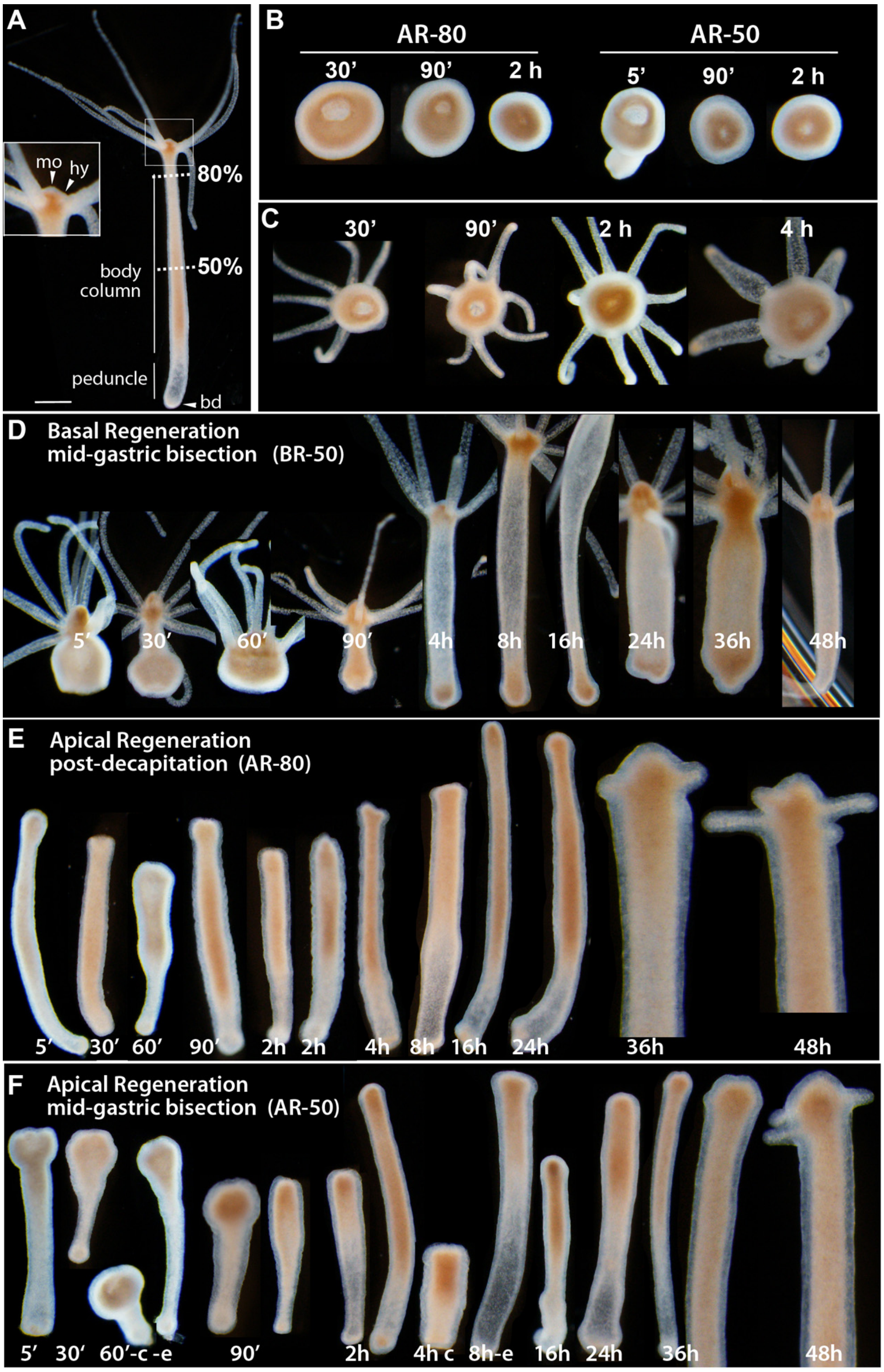

**Figure S 1. Anatomies of intact (A) or regenerating (B-F) *Hydra vulgaris* polyps after decapitation (80%, B, E) or mid-gastric bisection (50%, B, C, F)**

(**A**) Anatomy of an intact *Hydra*. bd: basal disc; hy: hypostome; mo: mouth opening. 80% and 50% indicate the decapitation and mid-gastric bisection levels respectively. Scale bar = 200 µm. (**B, C**) Transversal views showing the closure of the gastric wound during apical (B) and basal (C) regeneration. (**D**) Basal regeneration after mid-gastric bisection (BR-50). Note that the basal disc is formed at 36 hours post-amputation (hpa) and the animal can attach to the bottom of the dish as shown at 48 hpa. (**E, F**) Apical regeneration after decapitation (AR-80) and after mid-gastric bisection (AR-50). Tentacle rudiments become visible by 36 hpa in AR-80, several hours later in AR-50. During the first 12 hours of the regeneration process, animals exhibit spontaneous regular contractions as shown with views of the same animal contracted (c) or extended (e).

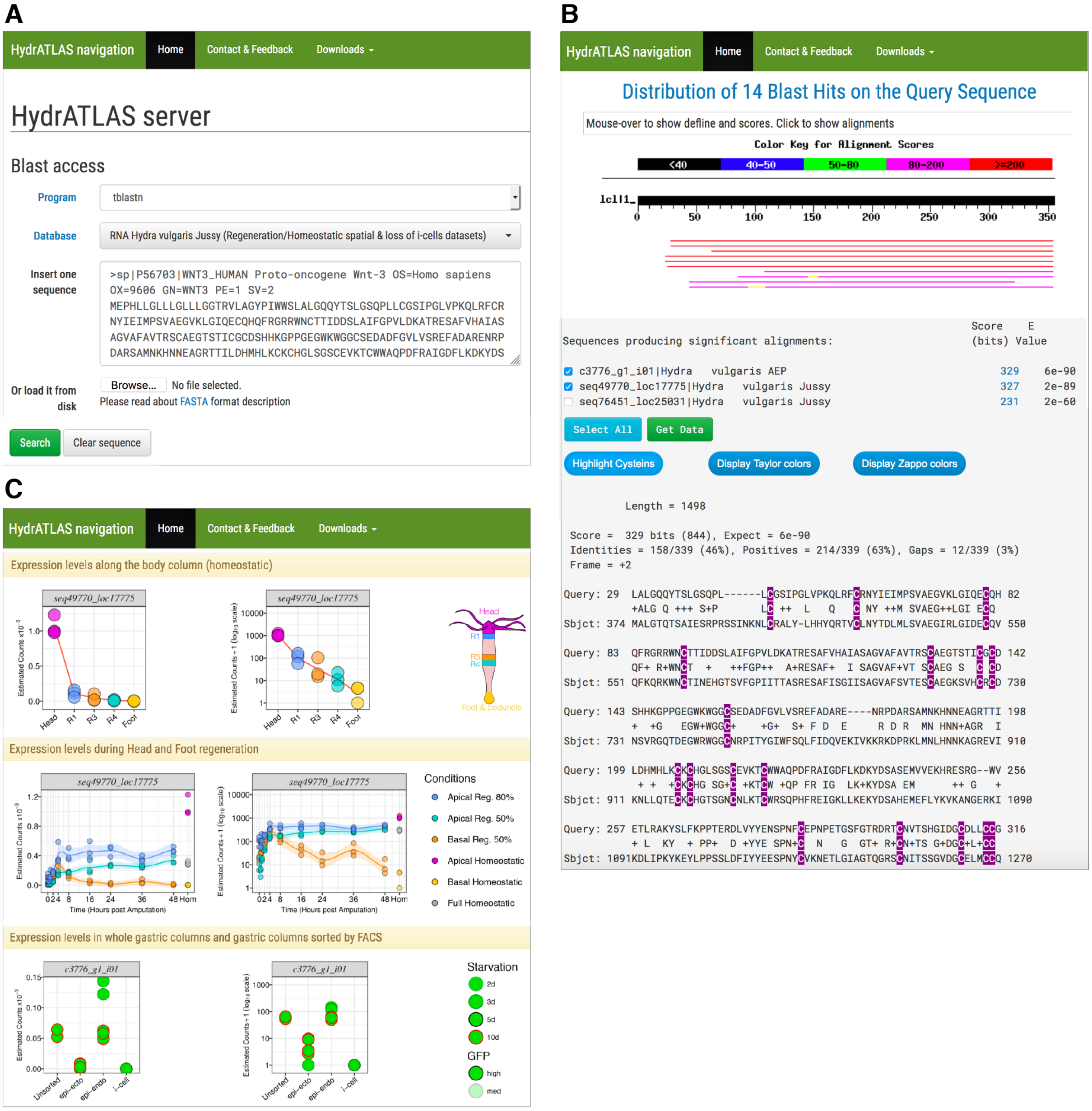

**Figure S 2. Schematic view of the HydrATLAS portal with the different contexts for RNA-seq profiling (https://hydratlas.unige.ch)**
**(A)** Queries can be fasta sequences from any organism or by *Hydra* sequence names (not shown). **(B)** blast alignment **(C)** Graphical output

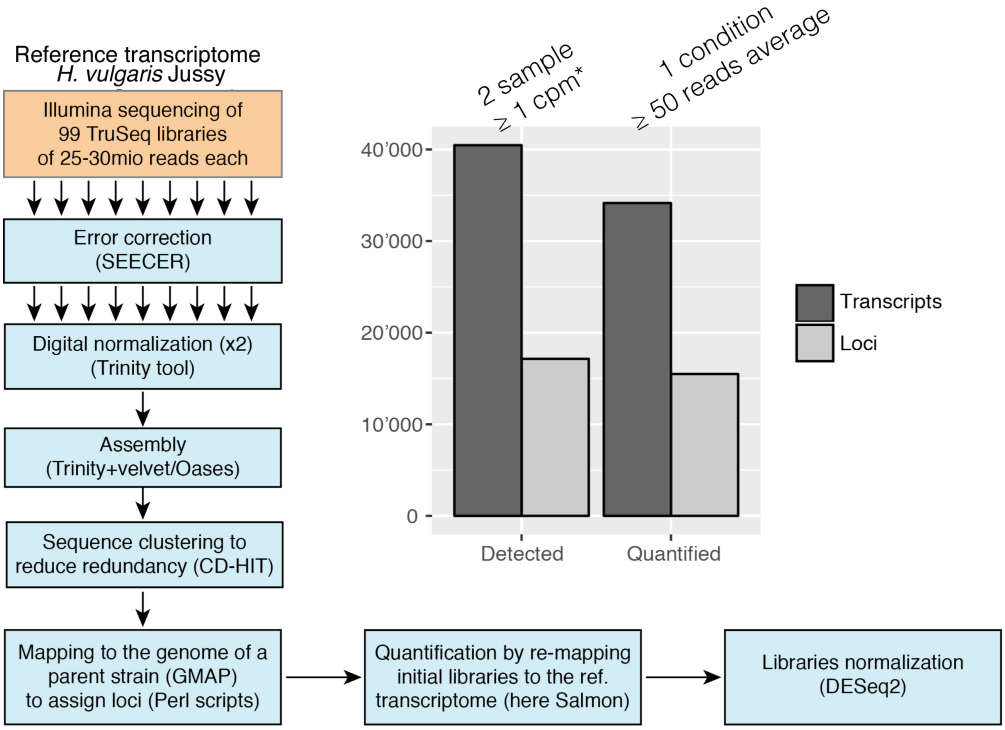

**Figure S 3. Technical conditions applied on RNA-seq Illumina reads to build the reference transcriptome of *H. vulgaris* Jussy strain**

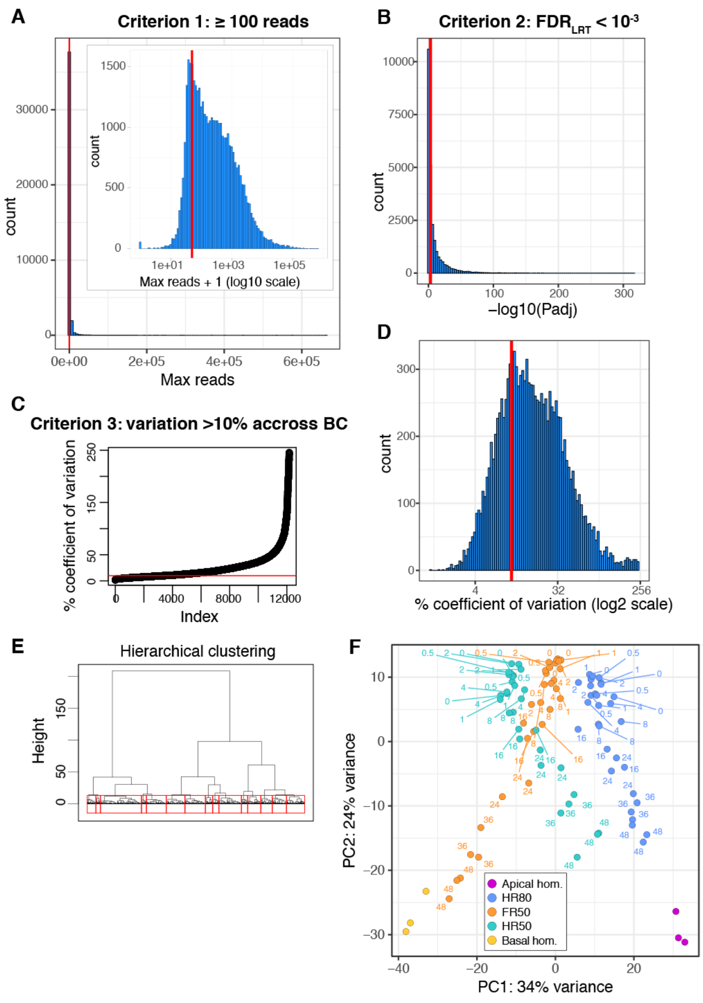

**Figure S 4. Criteria used for selecting transcripts submitted to the analysis of gene regulations in homeostatic and regenerative conditions**

BC: body column; FDR: False Discovery Rate

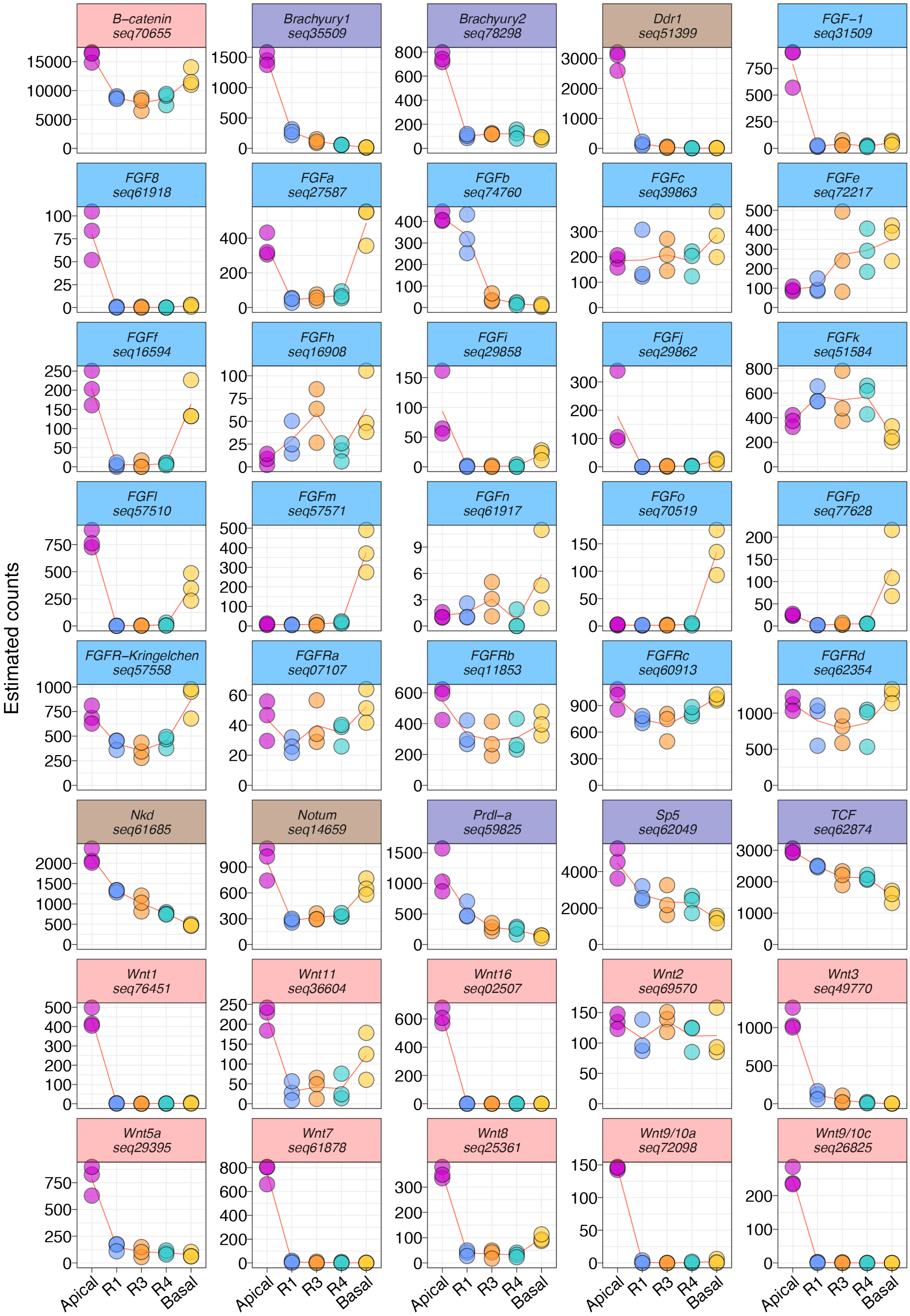

Figure S 5. Individual RNA-seq profiles of apical markers, basal markers, and transcripts related to Wnt and FGF signaling (related to Fig. 2A)

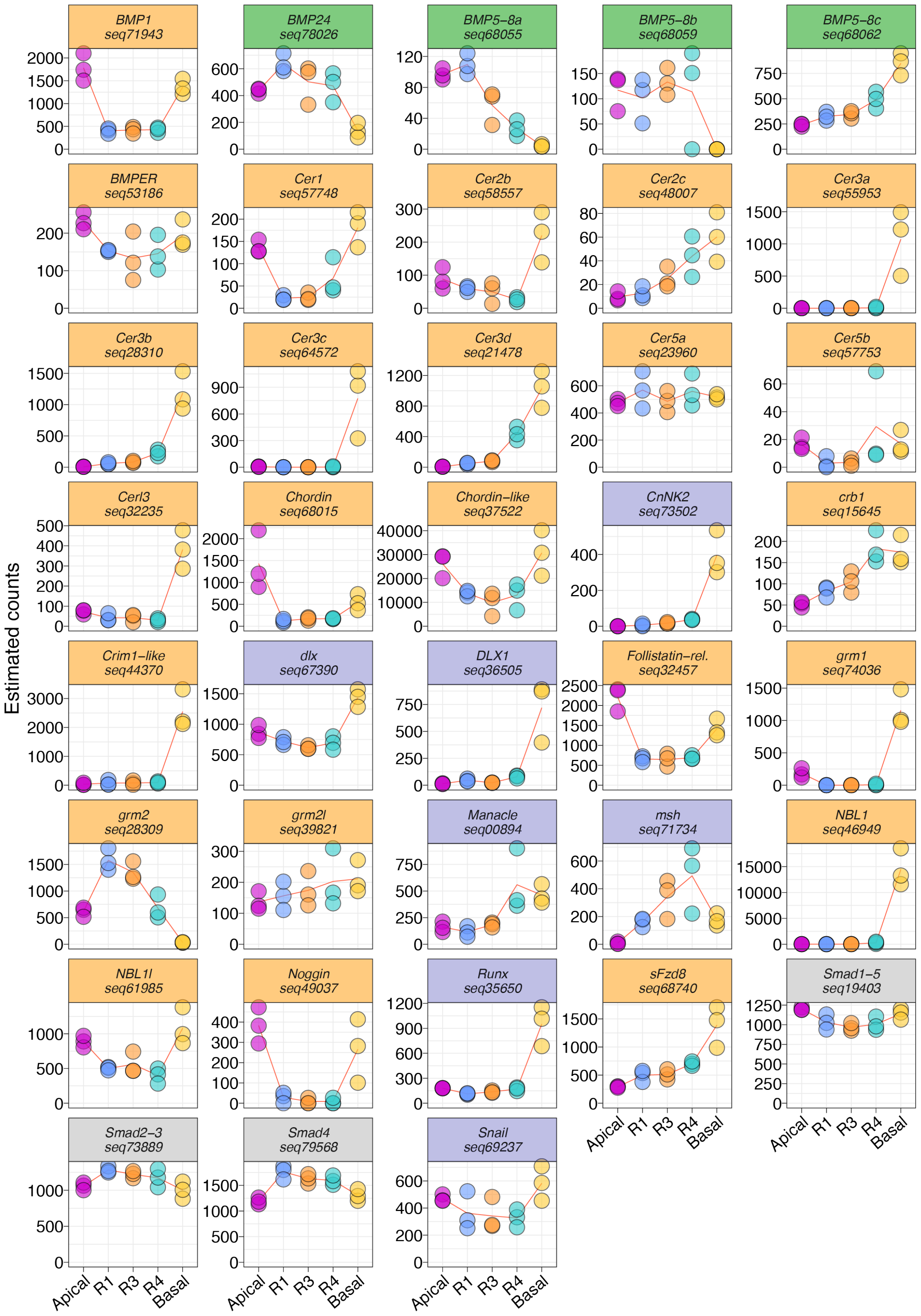

Figure S 6. Individual RNA-seq profiles of apical and basal markers, as well as transcripts related to BMP signaling (related to Fig. 2B)

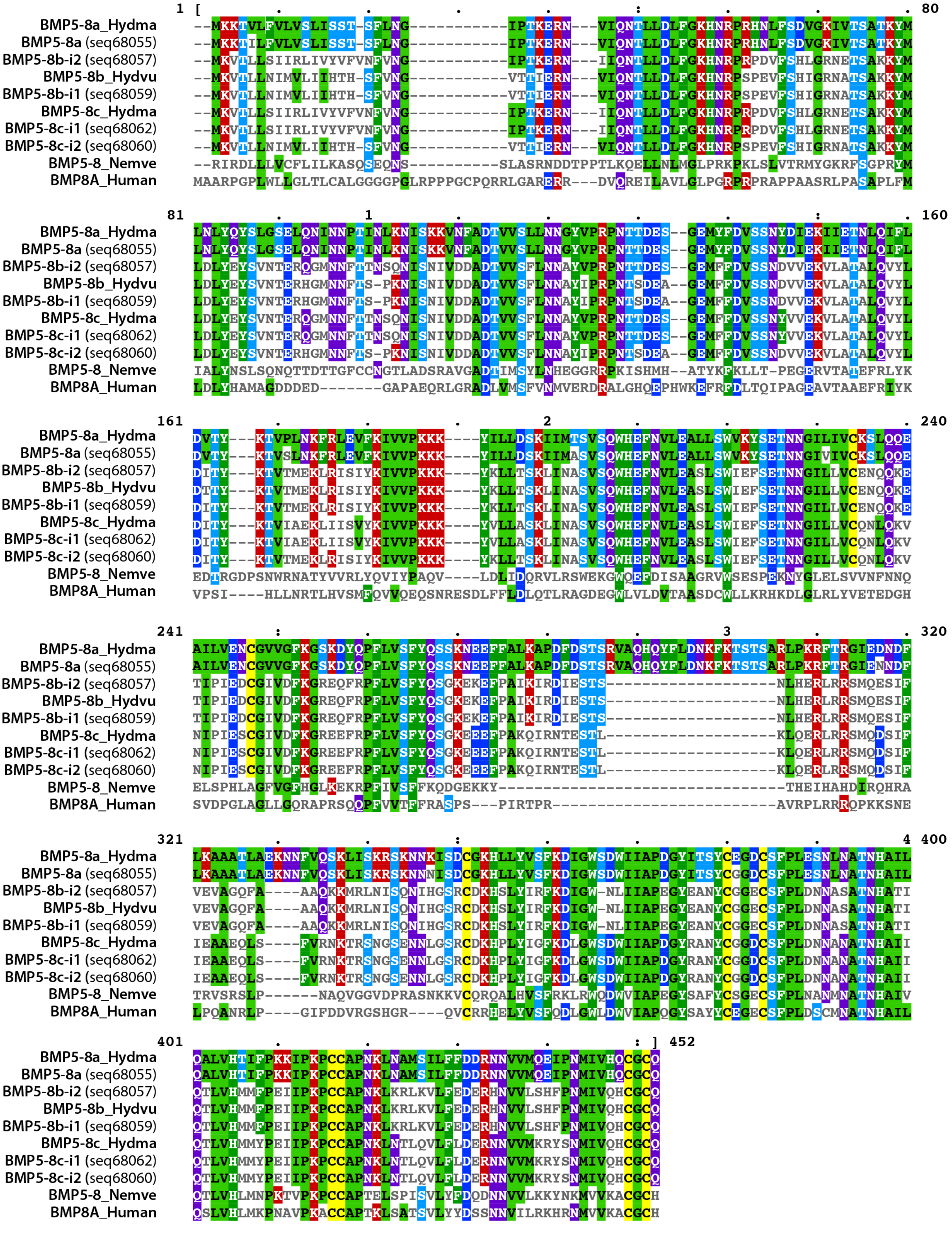

**Figure S 7. Multiple protein alignment of the *Hydra* BMP5-8 related sequences.**

Hydma: *Hydra magnipapillata*; seq68055, seq68057, seq68059, seq68060, seq68062 are deduced from transcripts isolated from *H. vulgaris Jussy* strain; Nemve: *Nematostella vectensis* (sea anemone).

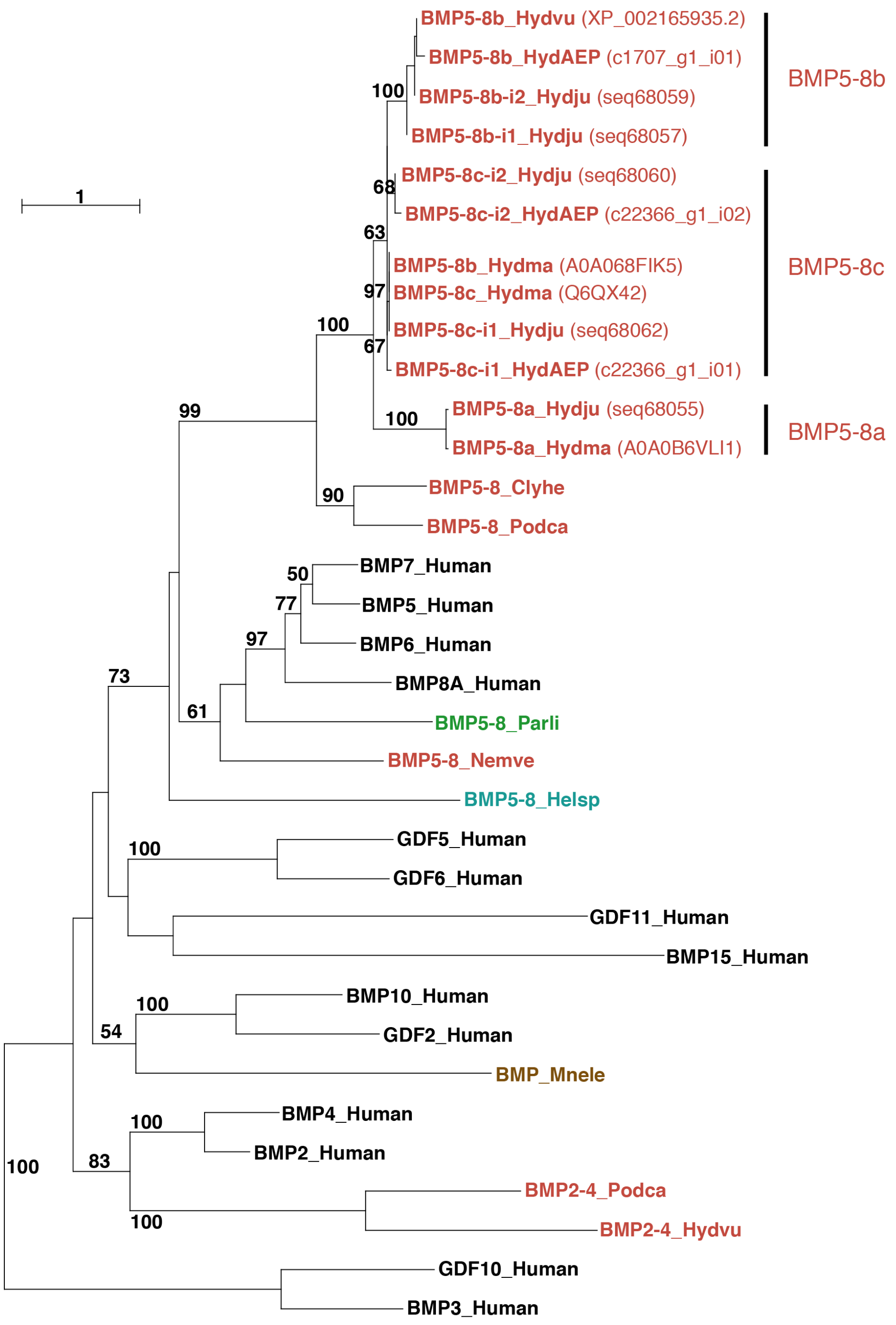

**Figure S 8. Phylogenetic analysis of *Hydra* BMP-related protein families**

*Hydra* BMP-related sequences identified from the *Jussy (HydJu)* and *AEP (HydAEP)* strains in the present RNA-seq transcriptomic analysis were compared to the BMP-related sequences available on Uniprot from *Clytia hemisphaerica (Clyhe*, hydrozoan jellyfish, A0A0A8P5M4), *Helobdella sp (Helsp,* leech, G1EH57*), H. magnipapillata (Hydma,* Q6QX42, A0A068FIK5, A0A0B6VNE3, A0A0B6VLI1*), H. vulgaris* (*Hydvu,* NCBI: XP_012554724.1), *Human* (P12643, P12645, P55107, P12644, P22003, P22004, P18075, Q7Z5Y6, Q9UK05, O95393, O95390, Q6KF10, P43026, O95972)*, Mnemiopsis leidyi (Mnele,* comb jelly – ctenophore, G5CTK5*), Nematostella vectensis (Nemve,* sea anemone, Q27W10*), Paracentrotus lividus (Parli,* sea urchin, A0A0N9R4G2), *Podocoryne carnea (Podca,* hydrozoan jellyfish, Q0R0B0, Q0R0B1*).* Color code of the protein sequences: cnidarian (red-orange), ctenophore (brown), echinoderm (green), annelid (blue), human (black). Sequences were aligned with Muscle Align (www.ebi.ac.uk/Tools/msa/muscle/) and the phylogeny was analyzed with Phyml 3.0 (Guindon et al., 2010) with LG as substitution model, 4 categories of substitution rates, estimated proportions of invariables sites, estimated gamma shape parameters and 100 bootstraps at the ATGC-Montpellier platform (http://www.atgc-montpellier.fr). Note the presence of three BMP5-8 paralogs in *Hydra* (see sequence alignment in Fig. S7).

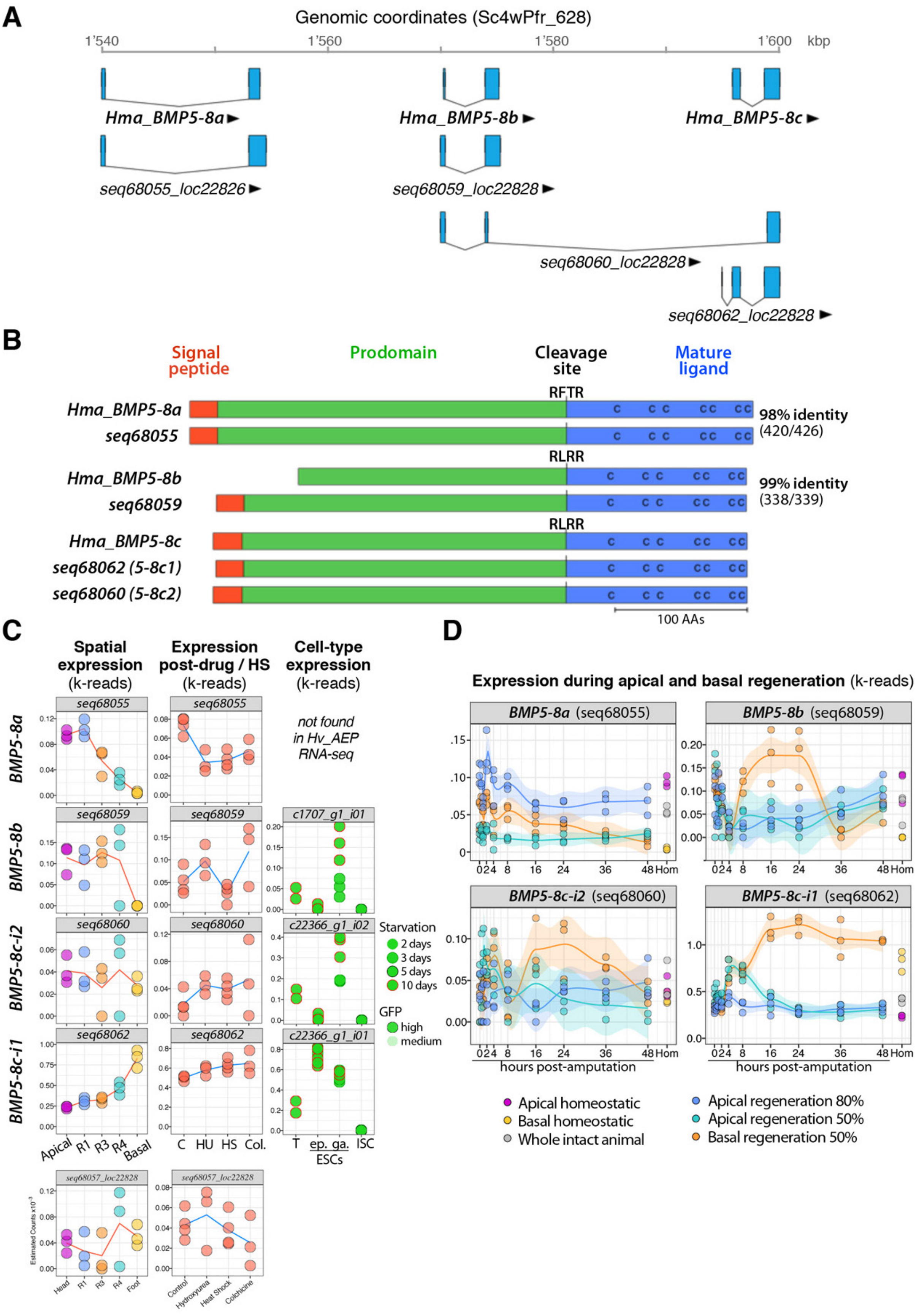

**Figure S 9. Genomic map and expression profiles of the three *Hydra* Bmp5-8 related genes**

**(A)** Genome alignment of the *Hydra Bmp5-8* transcripts containing the Transforming growth factor beta like domain (Pfam: PF00019). Bmp5-8 sequences from *H. magnipapillata* (*HmaBmp5-8a, HmaBmp5-8b, HmaBmp5-8c*) were retrieved from (Watanabe et al., 2014). Bmp5-8 sequences from *H. vulgaris Jussy* (seq68055, seq68059, seq68060, seq68062) were generated in the present study. Black arrows indicate 5' to 3' direction. The Hydra 2.0 genome assembly was used as reference. (https://research.nhgri.nih.gov/hydra/miscellaneous/about.shtml) **(B)** Schematic representations of the predicted Bmp5-8 proteins. All sequences generated by RNA-seq contain a signal peptide located at amino-acids positions 1 to 21/22, an extra-cellular prodomain ending with a dibasic cleavage site (-RXXR-) followed by a carboxy-terminal mature ligand. The mature ligand assumes a cystin-knot configuration with seven highly-conserved cysteins forming three disulfides bridges. (**C**) Homeostatic expression measured by RNA-seq at five positions along the body column (spatial expression), after elimination of the cycling interstitial cells by drug or heat-shock treatments, cell-type expression after FACS sorting of GFP expressing cells (transgenic Hydra AEP). eESC: epidermal epithelial stem cell, gESC gastrodermal epithelial stem cell, ISC: interstitial stem cell. HU: hydroxyurea, HS: heat-shock, colch.: colchicine. (**D**) Kinetics of expression during apical or basal regeneration after decapitation (80%) or mid-gastric bisection (50%).

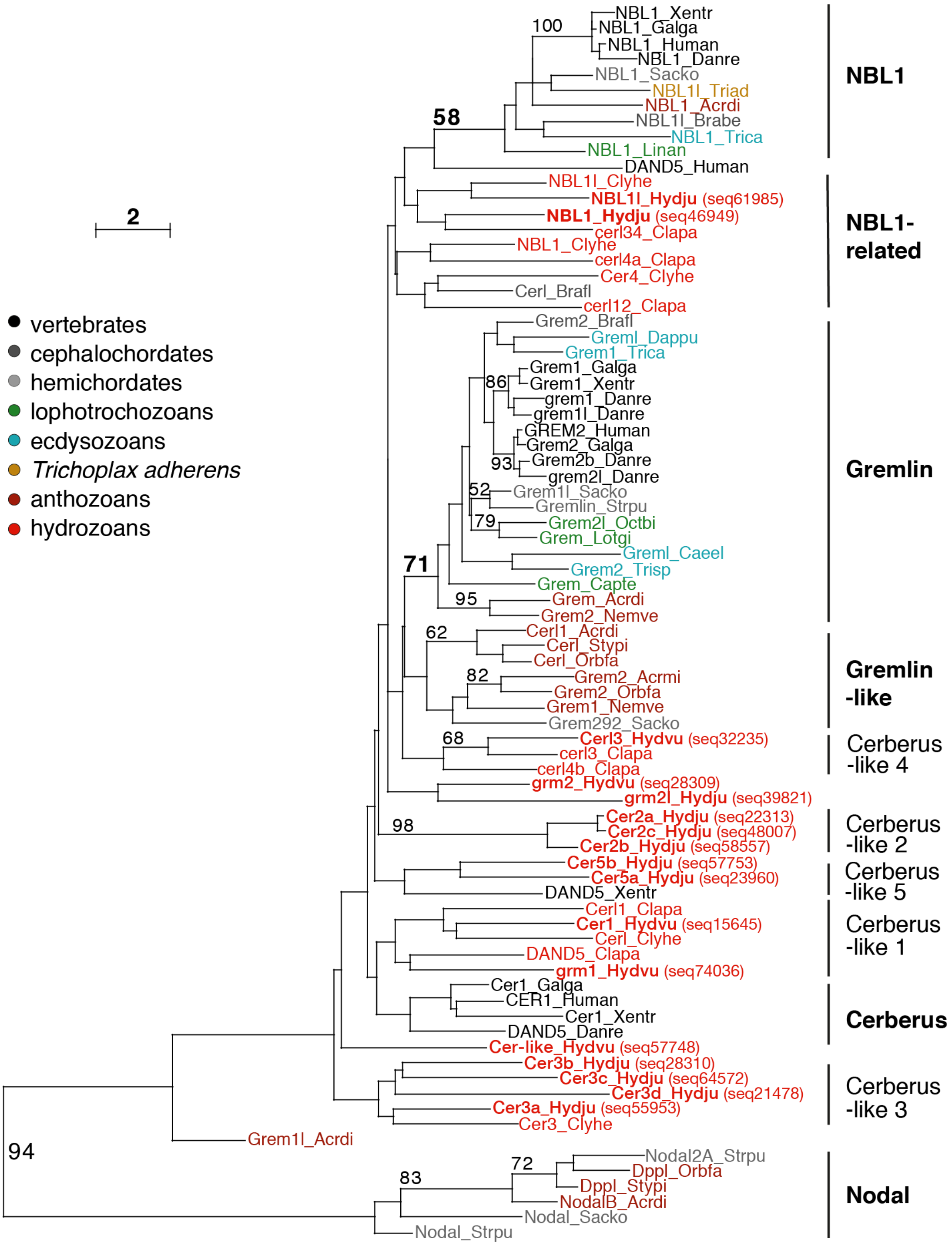

**Figure S 10. Phylogenetic analysis of the *Hydra* DAN, Gremlin and Cerberus protein families**

*Hydra* DNA-domain containing protein sequences were retrieved from the *Hv_Jussy* reference transcriptome and from (Watanabe et al., 2014). These sequences were then blasted on Uniprot and NCBI databases to be verified, annotated and completed with related sequences from cnidarian and bilaterian species. Related sequences from porifera or non-metazoan species were not found. 80 sequences were aligned with Muscle Align (www.ebi.ac.uk/Tools/msa/muscle/) and the phylogeny was analyzed with Phyml 3.0 (Guindon et al., 2010) with the LG substitution model, 8 categories of substitution rates, estimated proportions of invariables sites and gamma shape parameters. 100 bootstraps at the ATGC-Montpellier platform (http://www.atgc-montpellier.fr). Note the two sequences Cer-5b (seq57753) and NBL1 (seq46949) present in the genome but not characterized previously.

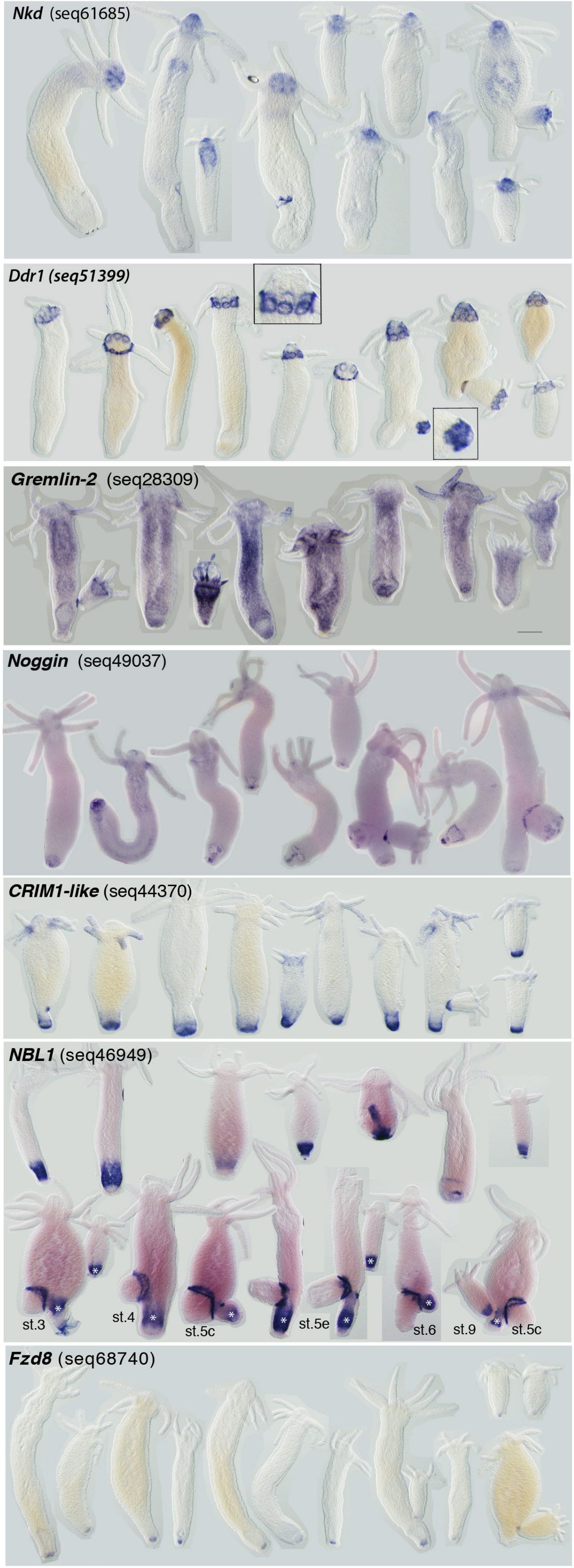

**Figure S 11. Expression patterns of candidate organizer regulators along the body axis of intact animals (*Hv_Basel*).**

The patterns detected by ISH confirm the spatial RNA-seq profiles with *Ddr1* apical, *Nkd, Gremlin-2* graded apical to basal, *Noggin* bipolar and *CRIM-1, NBL1, Fzd8* basal. See corresponding RNA-seq profiles in **Figure 3**.

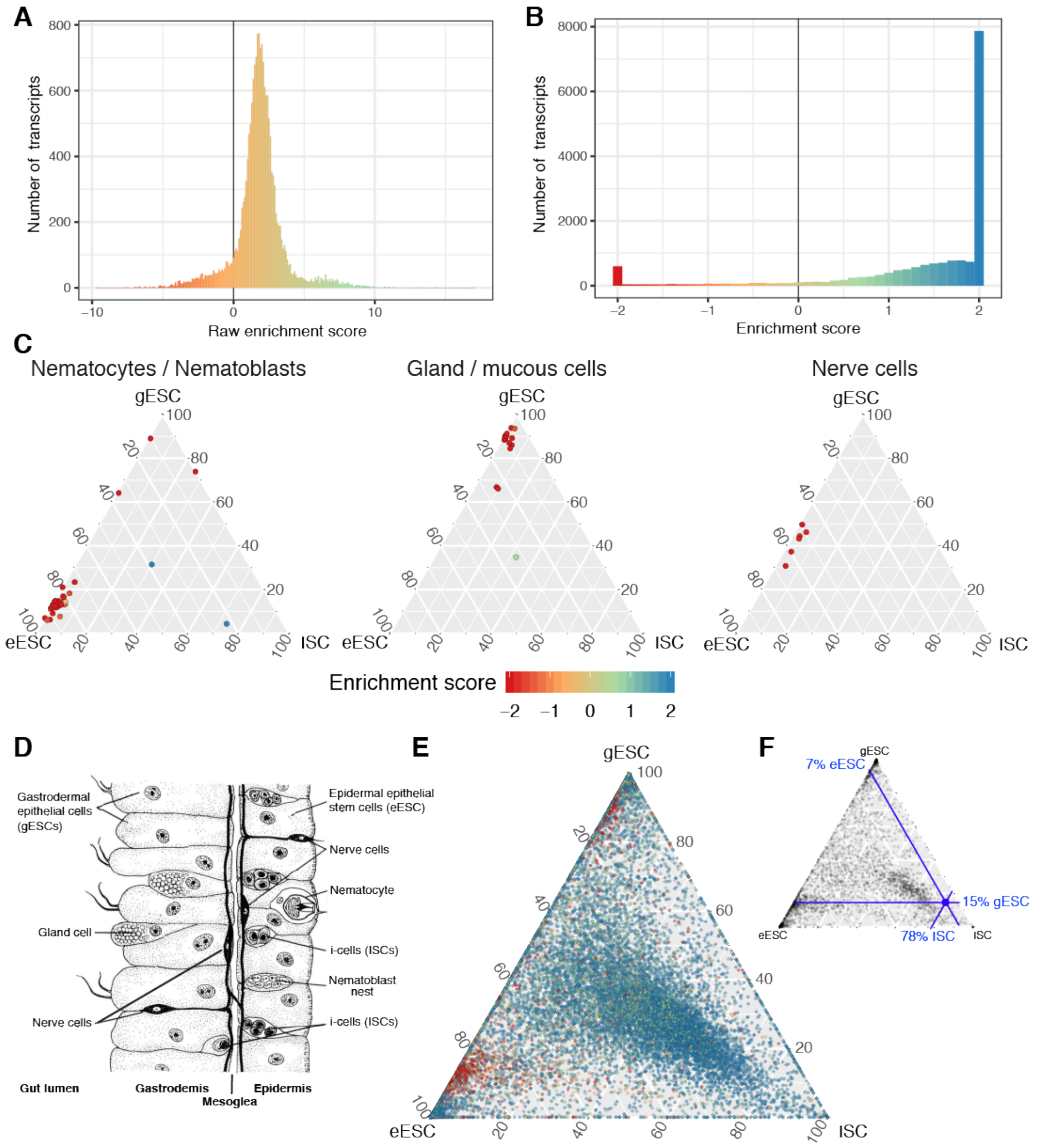

**Figure S 12. Enrichment score and identification of transcripts associated with ISC derivatives in the cell-type specific RNA-seq dataset**

**(A)** Raw enrichment scores for all transcripts identified in the dataset, showing that the vast majority of transcripts are enriched in the targeted cell types. **(B)** In order to produce a more detailed color scale, raw enrichment scores were constrained between -2 and +2, i.e. all values below or above than -2 or +2 were assigned the value -2 or +2. **(C)** RNA-seq Identification of transcripts from i-cell derivatives from a curated dataset of transcripts expressed in nematocytes/nematoblasts, gland/mucous cells, or nerve cells (Hwang et al., 2007). The position of each dot indicates its relative proportion among eESC, gESC, and ISC samples concentrated by FACS. Transcripts from nematocytes/nematoblasts-specific, a cell type located in the epidermal layer of the animal, were found predominantly in the eESC (epidermal epithelial) fraction. Gland cells in the gESC (gastrodermal epithelial) fraction, and nerve cells in roughly equal amounts in the gESC and eESC fractions. As the majority of these transcripts were found at much elevated levels in the unsorted gastric tissue, most of their enrichment scores are low (red color), allowing to discriminate them from transcripts expressed in the three targeted stem cell populations. **(D)** Localization of cells in *Hydra* tissue layers. Image adapted from (Holstein and Emschermann, 1995). **(E)** Cell type specific distribution could be attributed to 17'561 transcripts using an expression threshold of more than 100 reads in one or more of the samples. **(F)** The position of the i-cell specific transcript *nanos/CnNos1* (blue dot, c19803_g1_i01, enrichment score: 2+) indicates that i-cells transcripts are highly enriched in the ISC fraction (78% of all counts), but that they are also present in gESC (14%) and to some extent in eESC (6%). The set of transcripts represented in **(E)** and **(F)** is identical, but coloring by enrichment score and scales have been omitted for readability.

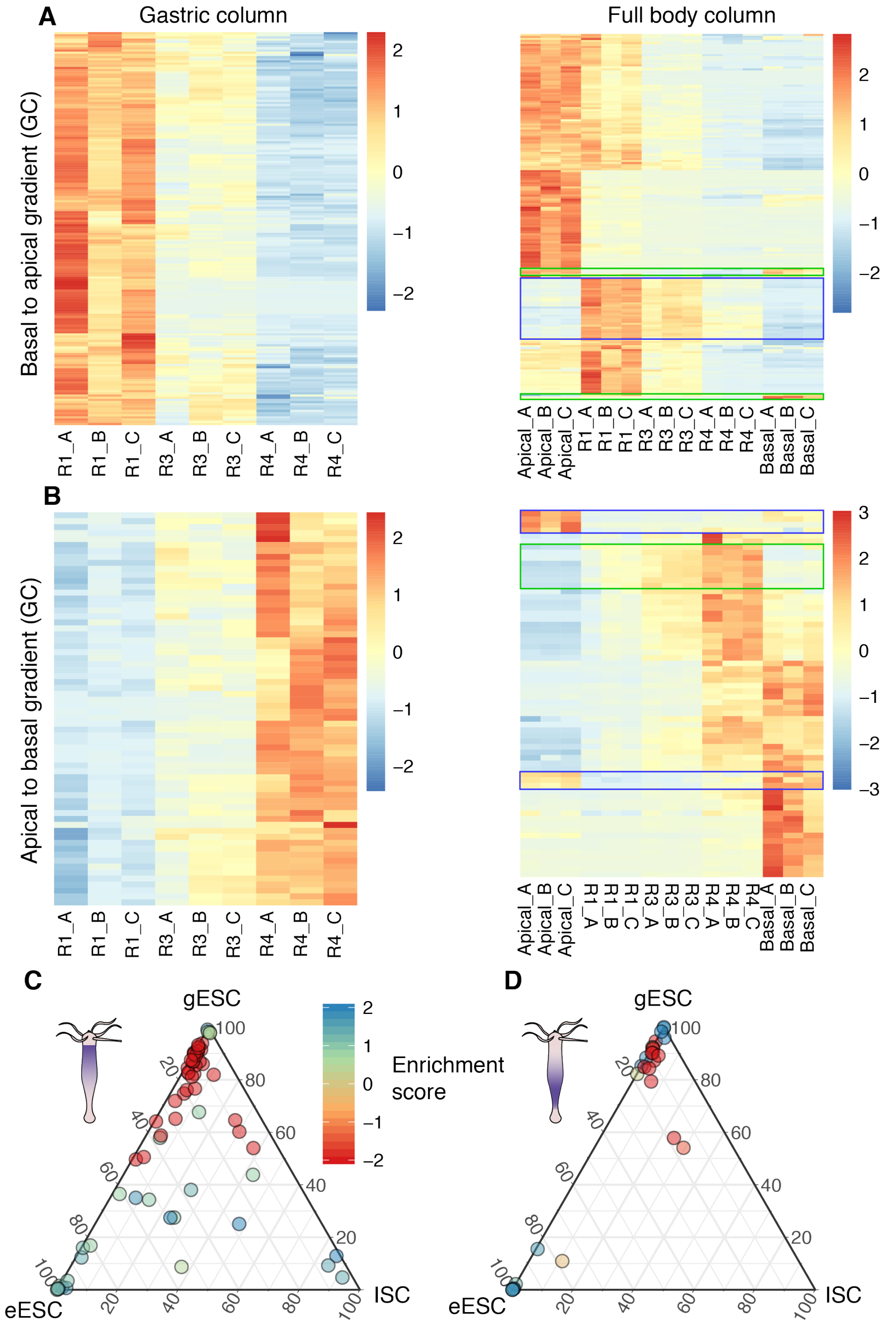

**Figure S 13. Transcripts with strongly graded expression in the gastric region are predominantly expressed in gland cells.**

Transcripts with strongly graded expression along the gastric regions were selected as follows (for basal to apical graded transcripts): the minimum value recorded in the R1 region is higher than the maximum value of R3 triplicates. Similarly, values in R3 triplicates are all higher than R4 triplicates. A similar procedure was applied to identify apical to basal transcripts. Note that expressions in the basal and apical regions were not considered to identify transcripts graded along the gastric region **(A)** (left) Heatmap representation of 180 transcripts with graded basal to apical expression in the gastric region, (right) extension of the previous heatmap to represent apical and basal regions. Data are library-normalized estimated counts standardized by transcript. **(B)** 66 transcripts satisfying the criteria for graded apical to basal expression in the gastric region. **(C)** ternary plots representing the cell type expression of the transcripts identified as graded basal to apical (shown in (A)). **(D)** ternary plots representing the cell type expression of the transcripts identified as graded apical to basal (shown in (B)). After mapping *Hv_Jussy* transcripts to the *Hv_AEP* transcriptome by reciprocal best hits, 31 and 100 transcripts with cell type information remained for apical to basal expression and basal to apical expression, respectively. Blue and green boxes show transcripts with independent modulations at the extremities for the apical region and for the basal region, respectively.

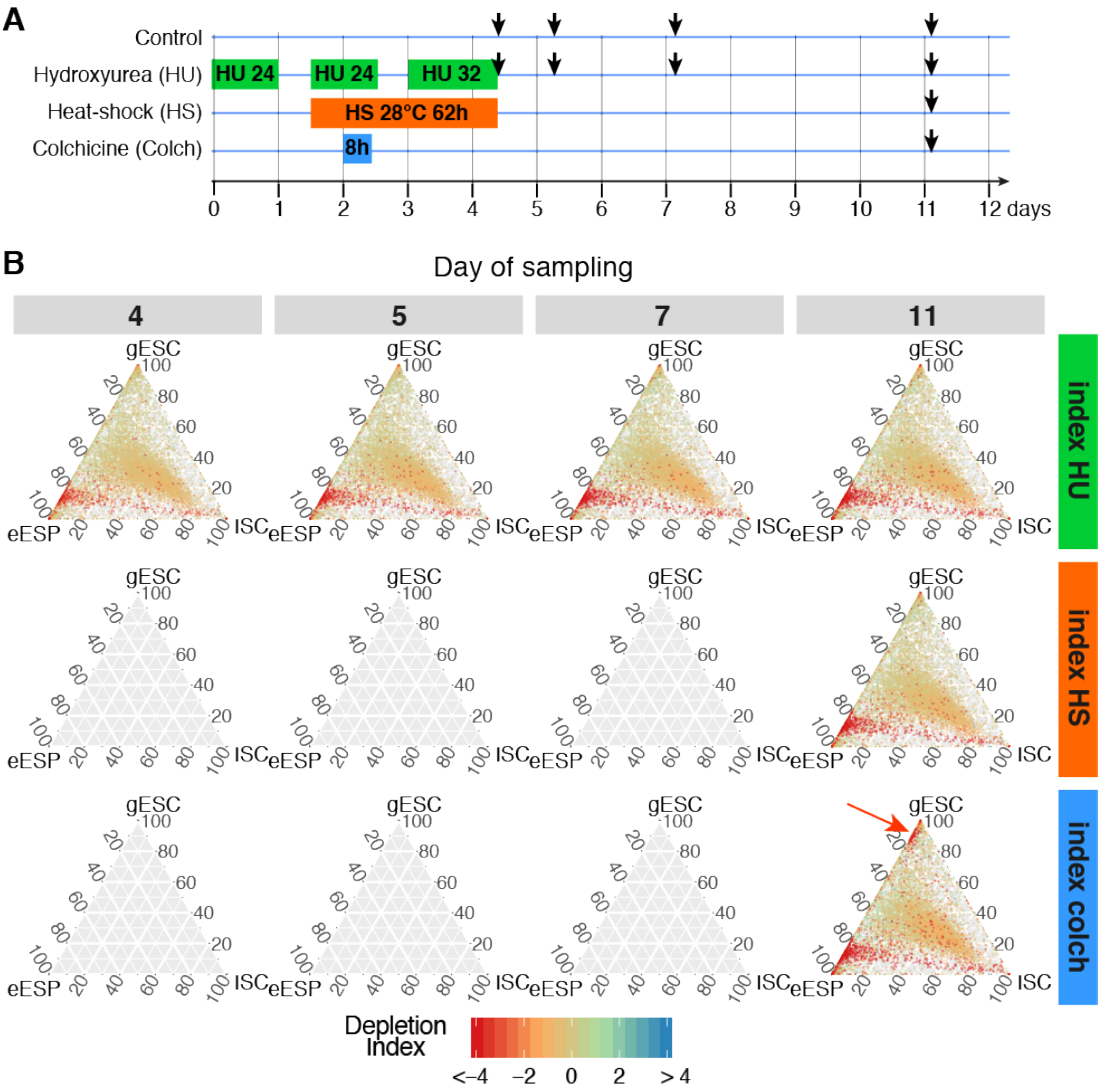

**Figure S 14. Kinetics of transcript depletion upon interstitial cell loss induced by treatments targeting fast-cycling cells**

(**A**) Experimental design of the hydroxyurea (HU), heat-shock (HS) and colchicine (Colch) experiments leading to the elimination of the fast cycling cells in *Hydra*, i.e. ISCs and interstitial progenitors collectively named i-cells (i-cell loss). Black arrows indicate the days when the central body columns of treated or untreated animals were dissected for RNA preparation. Experiments were performed on *H. vulgaris sf-1* strain as depicted in Wenger et al. (2016). (**B**) Ternary plots representing the depletion of transcripts after treatments to HU/HS/Colch. The depletion index is calculated for each transcript as the log-ratio of its abundances between the treated and the control condition at a given day and for a given treatment. In contrast to the enrichment score presented in **Fig. 4B**, depletion indexes calculated here have value below zero for transcripts reduced by the treatment. The red arrow indicates the depletion in the gESC fraction, likely corresponding to transcripts expressed in gland cells sampled as contaminant of the gESC fraction. Loss of gland cell transcripts is only detected after colchicine treatment.

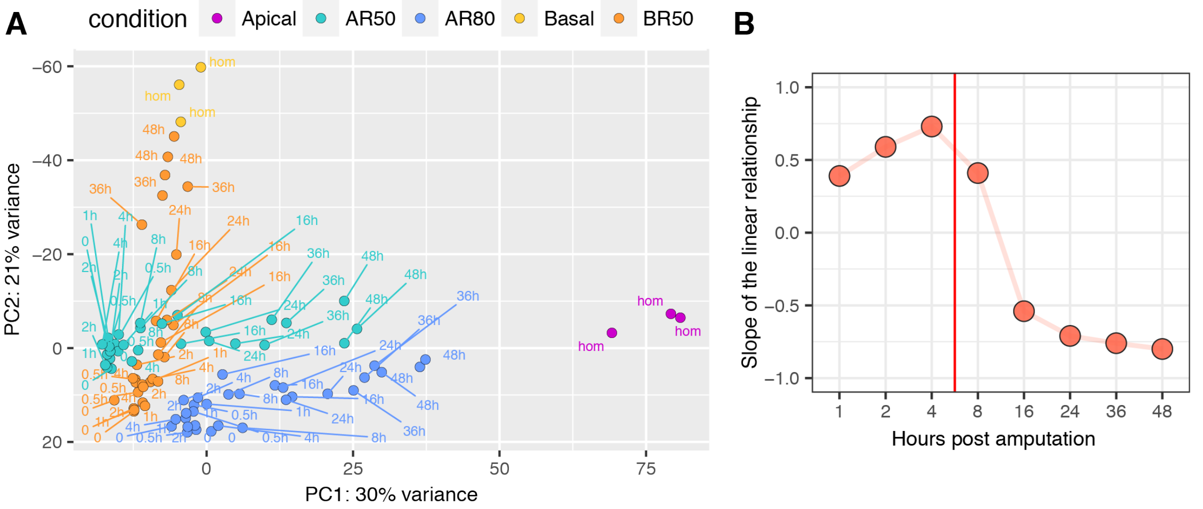

Figure S 15. PCA analysis of regenerating tips (with triplicate samples) and summary of slope variations over time (related to Fig. 5)

**(A)** Similar analysis than presented in Fig. 5B but including all triplicates. hom: homeostatic condition. **(B)** Summary plot of the slopes of the linear regressions presented in Fig. 5C.

---

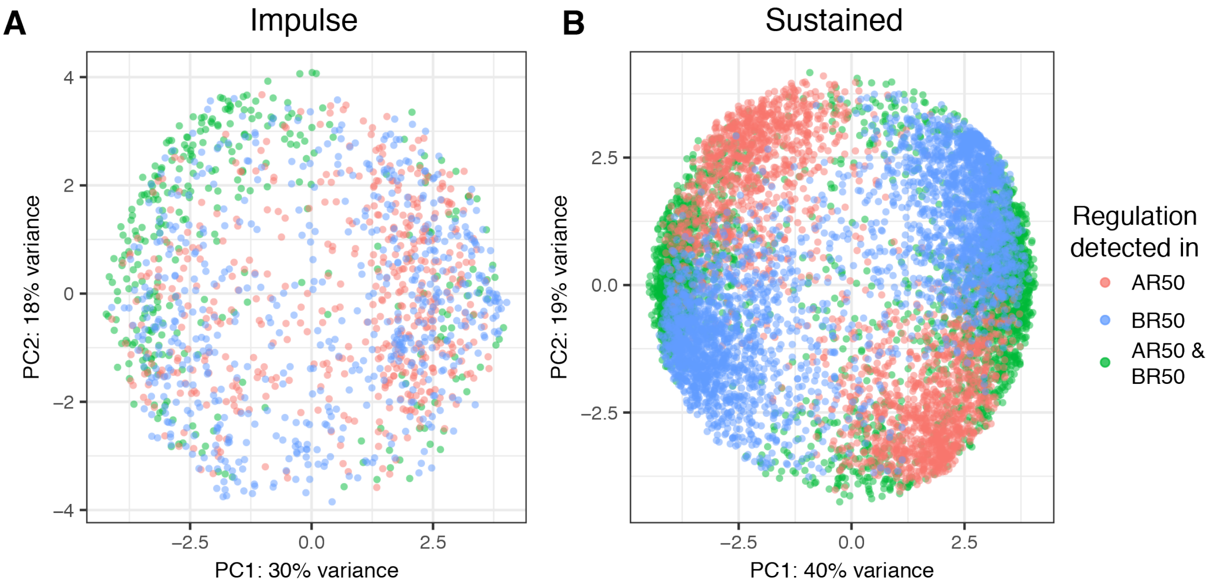

Figure S 16. PCA analysis of transcripts detected as impulse or sustained

PCA analyses performed on normalized counts of the regeneration time course after rlog transformation and scaling either on transcripts classified as **(A)** impulse or **(B)** sustained in one or both regenerative conditions. **(A)** The homogeneous distribution of transcripts detected as impulse (no obvious patterns or clusters for any of the conditions) indicates that expression patterns of transcripts in AR50 and BR50 conditions are not different for the majority of impulse-type transcripts. **(B)** Among transcripts with sustained expression profiles, some genes exhibit similar profiles in AR50 and BR50 but a majority are segregated.

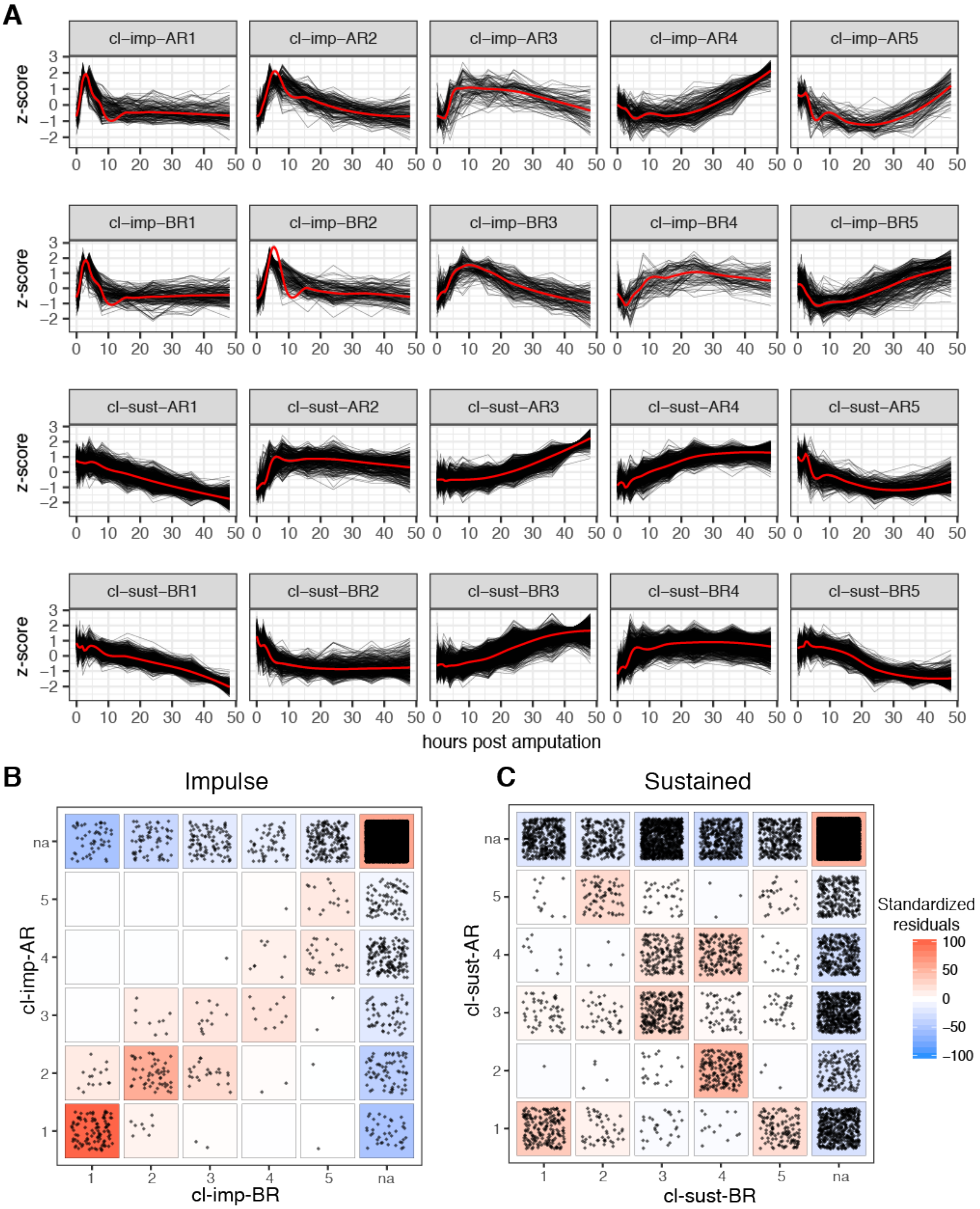

**Figure S 17. Transcripts detected as impulse or sustained grouped by k-means clustering**

**(A)** 683 transcripts with impulse regulation during apical regeneration (cl-imp-AR), 744 transcripts with impulse regulation during basal regeneration (cl-imp-BR), 3'231 transcripts with sustained regulation during basal regeneration (cl-sust-AR), 3'846 transcripts with sustained regulation during basal regeneration (cl-sust-BR). Black lines represent individual transcripts, red lines are loess models. Standardized values (z-scores) were calculated on normalized estimated counts after regularized log transformation. Scatterplot of transcripts identified with **(B)** impulse or **(C)** sustained regulations. Each dot represents a transcript. na: not attributed as either impulse or sustained. Color code: standardized residuals from chi-squared test, red indicate overrepresentation and blue underrepresentation.

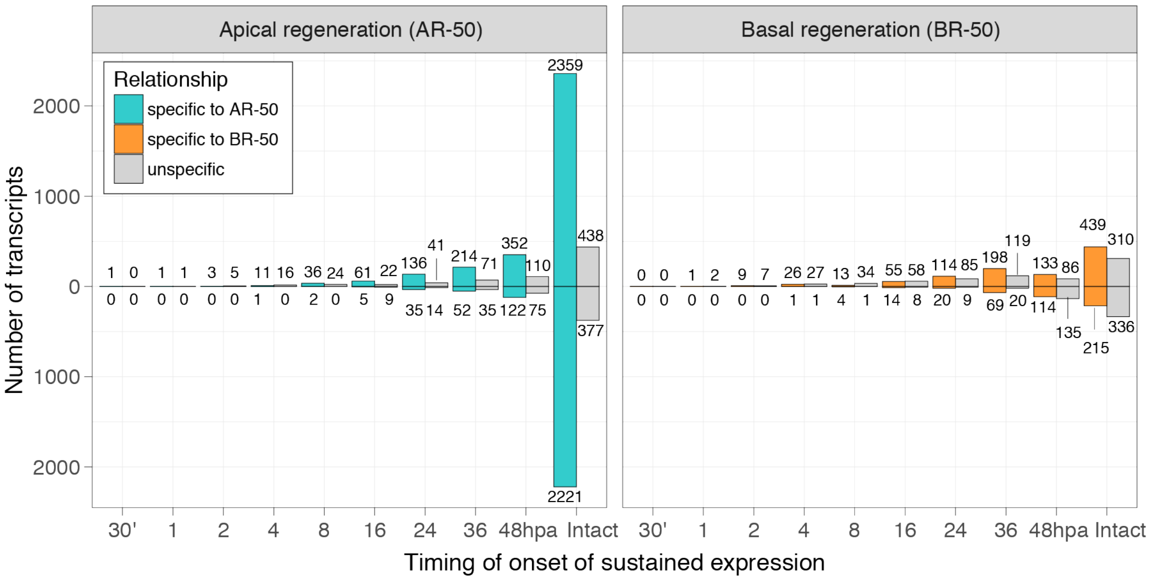

**Figure S 18. The number of transcripts up-regulated in a sustained manner increases over time**

**(A)** The onset is the first time point when a regulation (positive or negative, compared to time 0) is recorded and is maintained qualitatively at all subsequent timepoints and in the final structure to be regenerated (FC > ±1.5x, FDR < $10^{-3}$). The initial condition is region R4 for AR-50 and region R3 for BR-50 (**Fig. 1A, 2A**). Numbers in the graph correspond to initial timings of onsets. Bold letters (B to G) indicate at which time point initial timings of onsets are detected for transcripts presented in lower panels (B-G). Only a few genes have a sustained onset before 4 hpa and the number of sustained onsets increases over time, except for up-regulated transcripts in the BR-50 condition at 48 hpa.

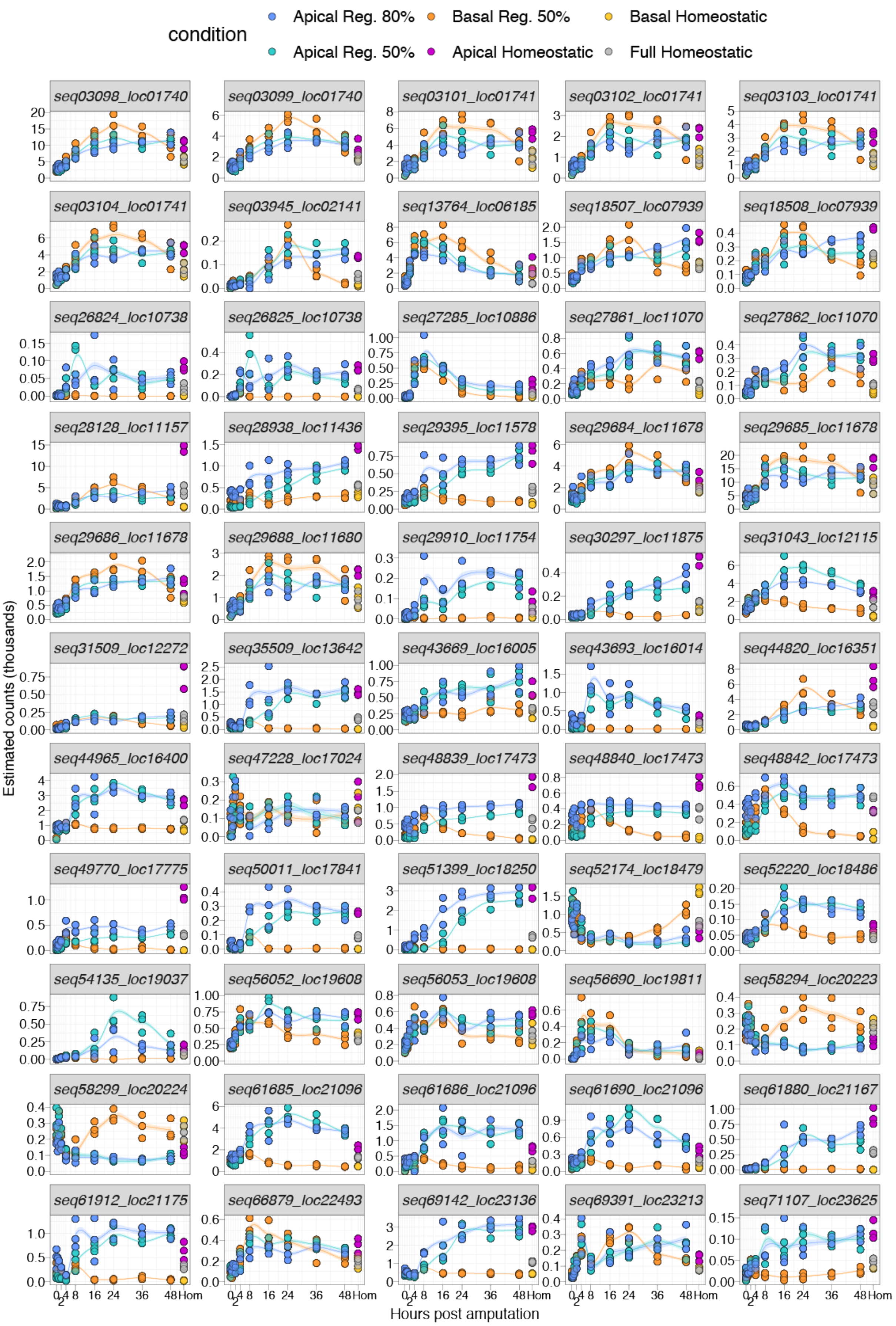

**Figure S 19. Transcripts showing a sustained up-regulation initiated within the first 8 hours after bisection in the head-regenerating tips**

(corresponding to Fig. 7A, left panel)

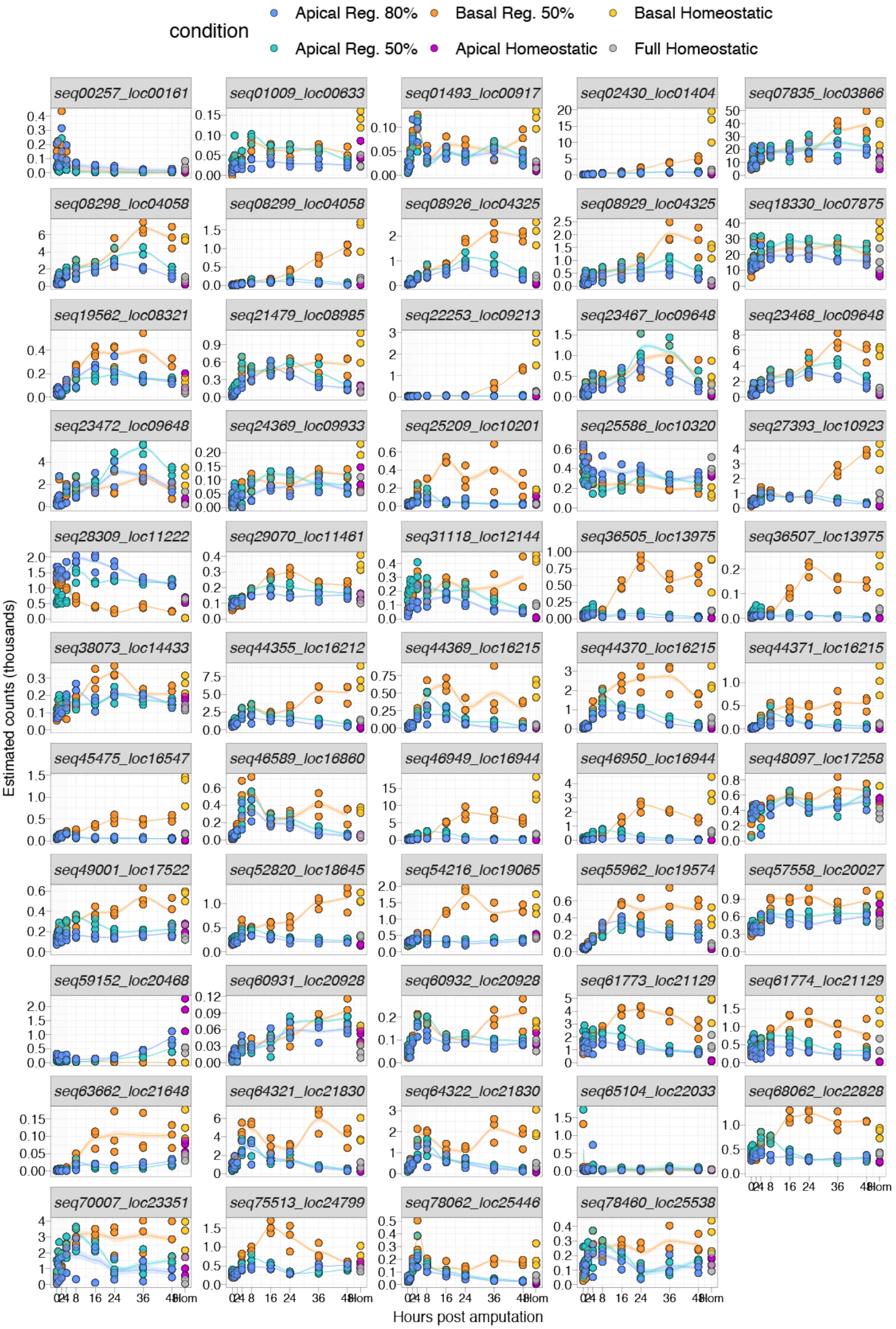

**Figure S 20. Transcripts showing a sustained up-regulation initiated within the first 8 hours after bisection in the basal-regenerating tips**

(corresponding to Fig. 7A, right panel)

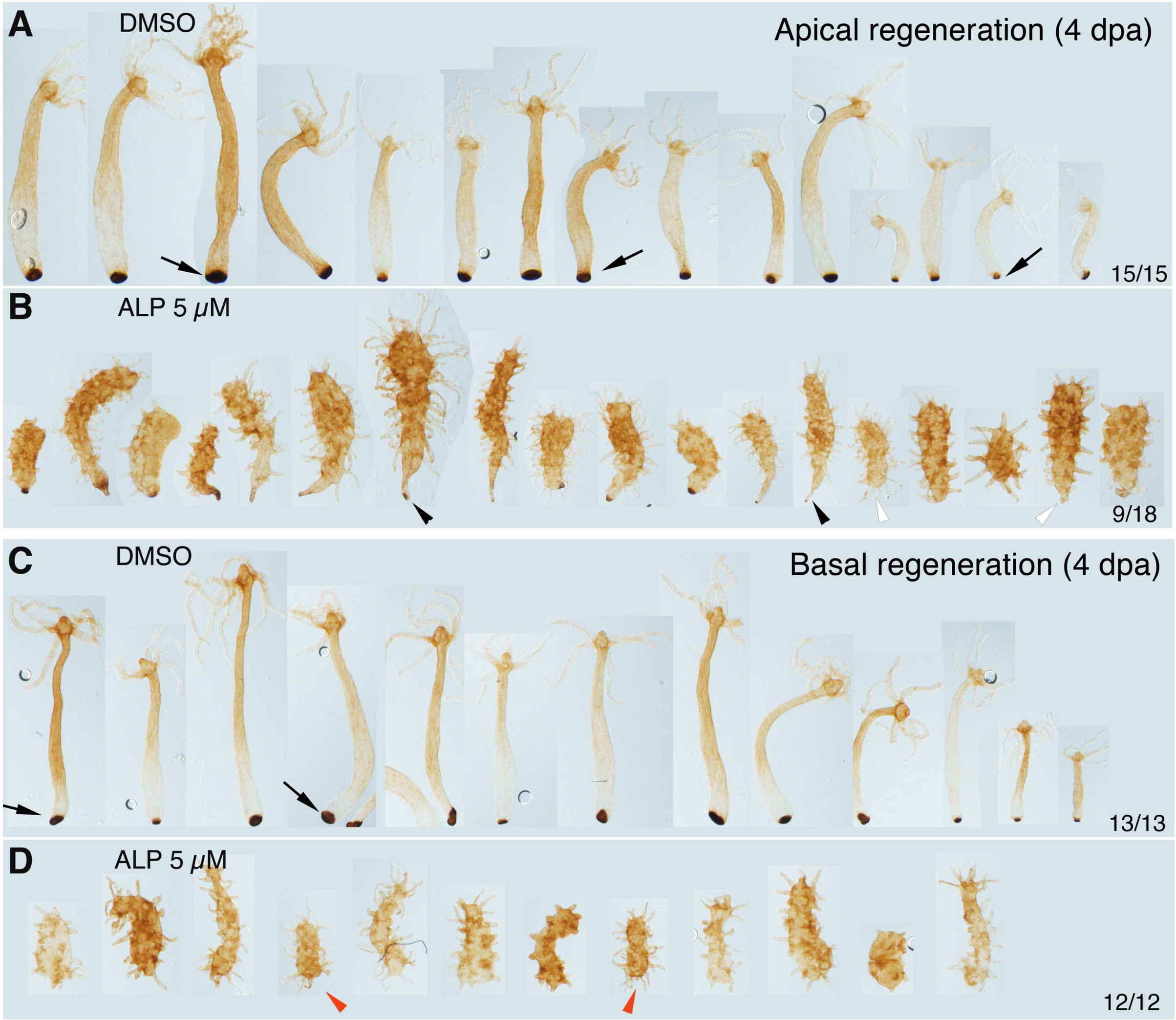

**Figure S 21. Loss of basal differentiation and basal regeneration in ALP-treated animals**

Peroxidase detection 4 days after mid-gastric bisection in halves having regenerated apical **(A, B)** or basal **(C, D)**. Animals in B and D were exposed to ALP for two days prior to bisection, four days later all of them display ectopic tentacles all along the body column. **(A)** All untreated animals have regenerated their apical half and maintained their basal disk intact as shown by the strong peroxidase activity (arrows, 15/15). **(B)** In ALP-treated-animals apical regeneration is altered and the original basal discs are either drastically reduced (black arrowheads) or have completely disappeared (white arrowheads, 9/18). **(C)** All untreated animals have regenerated their lower half and differentiated a new basal disk as shown by the strong peroxidase activity (black arrows, 13/13). **(D)** In ALP-treated-animals basal regeneration is abolished as evidenced by the lack of peroxidase activity at the basal pole (red arrowheads, 12/12). (see corresponding Fig. 9G)
